# Simulation of the Gαq /Phospholipase Cβ1 Signaling Pathway Returns Differentiated PC12 Cells to a Stem-like State

**DOI:** 10.1101/2020.01.06.896530

**Authors:** Osama Garwain, Katherine M. Pearce, Lela Jackson, Samuel Carley, Barbara Rosati, Suzanne Scarlata

## Abstract

Phospholipase Cβ1 is activated by Gαq to generate calcium signals in response to hormones and neurotransmitters, and is found at high levels in mammalian neuronal tissue. Besides carrying out this key plasma membrane function, PLCβ1 has a cytosolic population that helps, in part, to drive the differentiation of PC12 cells by inhibiting a nuclease that promotes RNA-induced silencing (C3PO). Here, we show that down-regulating PLCβ1 or reducing its cytosolic population by activating Gαq to drive it to the plasma membrane, returns differentiated PC12 cells to an undifferentiated state. In this state, the cells return to a spherical morphology, resume proliferation and express the stem cell transcription factors nanog and Oct4. Similar changes are seen with C3PO down-regulation. This return to a stem-like state is accompanied by shifts in multiple miR populations, such as increased levels of rno-miR-21 and rno-miR-26a. Surprisingly, we find that de-differentiation can also be induced by extended stimulation of the Gαq. In this case, the neurites completely retract over a 10-minute period, and while levels of nanog remain unchanged, the levels of some miRs begin to return to their undifferentiated values. In complementary studies, we followed the real time hydrolysis of a fluorescent-tagged miR in cells where PLCβ1 or C3PO were down-regulated. These samples showed substantial differences in miR processing in cells both the undifferentiated and differentiated states. Taken together, our studies suggest that PLCβ1, through its ability to regulate C3PO and endogenous miR populations, plays a key role in mediating PC12 cell differentiation.

## INTRODUCTION

Differentiation of a stem cell to a dedicated lineage occurs through a series of carefully orchestrated transcriptional events that generate the transcription of specific mRNAs and alter the distribution and amounts of proteins in the cell. While many of these events occur in rapid succession, some occur over longer periods resulting in cells that may have a narrow or broad a distribution of differentiation. Reversal of differentiation often occurs in cells that have transformed to cancerous states [1]. Thus, the ability to control differentiation and induce de-differentiation would greatly aid in optimizing cancer therapies and other therapies where inducing or reversing differentiation would be beneficial.

In this study, we show that we can manipulate the differentiated state of PC12 cells, a cultured cell line that is commonly used as a model for neuronal cells, through the Gαq/PLCβ1 signaling pathway [2, 3]. This pathway is initiated when neurotransmitters such as acetylcholine, serotonin, dopamine and melatonin bind to their specific transmembrane receptor. The ligand-bound receptor then activates Gαq which in turn activates phospholipase Cβ, leading to an increase in intracellular calcium. There are four known families of PLCβ, and PLCβ1 is the most responsive to Gαq, and is highly expressed in neuronal cells, such as PC12.

Aside from its classic role on the plasma membrane, PLCβ1 also has an atypical cytosolic population that binds to unique partners. Specifically, we have found that PLCβ1 binds to and inhibits component 3 promoter of RISC activity (C3PO), which promotes RNA-induced silencing by accelerating degradation of the passenger strand of the silencing RNA [4, 5]. Binding to C3PO occurs through the non-catalytic C-terminal region of PLCβ1, which is required for binding and activation by Gαq. The lipase activity of PLCβ is not required for C3PO inhibition, and C3PO does not inhibit PLCβ. However, C3PO competes for the same site on PLCβ as Gαq and increasing its cellular levels quenches calcium signaling mediated through Gαq receptors [6–8]. It is notable that increasing the level of activated Gαq by transfection or stimulation, will result in dissociation of PLCβ from C3PO and allow C3PO to promote RNA-induced silencing [7, 8].

In previous studies, we found that cytosolic PLCβ1 is required for differentiation of PC12 cells [9]. PC12 cells are a widely used model for neuronal cell differentiation [10]. In their undifferentiated state, they are round, rapidly dividing cells. After exposure to the neurotropic factor NGF (nerve growth factor), they stop dividing, sprout neurites, and form and release synaptic vesicles within ∼72 hours. Although changes induced by NGF are initiated by TrkA receptors that signal though pathways associated with growth factors [11], we have shown that PLCβ1 and C3PO are absolutely necessary for the morphological changes that accompany differentiation [9]. Additionally, in early stages of differentiation, PLCβ1 binds and inhibits CDK16 [12], a testis and neuronal-specific cyclin dependent kinase that promotes proliferation. The bifunctional effects of PLCβ1 on differentiation can be reversed by increasing the level of activated Gαq to drive the cytosolic population of PLCβ1 associated with CDK16 and C3PO to the plasma membrane thereby returning cells to a round, proliferative state.

Since the level of Gαq impacts the level of cytosolic PLCβ1 which is required for PC12 cell differentiation, then external signals that activate Gαq may impact the state of differentiation. Here, we show that down-regulating PLCβ1 in fully differentiated PC12 cells returns cells to a stem-like state in terms of morphology, division and stem cell markers. Our data suggest that the mechanism responsible for de-differentiation is lies in the reduction of PLCβ1-C3PO complexes that affect specific cellular miR populations and in turn, expressed proteins. Prolonging the activated state of Gαq also returns PC12 cells to an undifferentiated state. Our results suggest that PLCβ1 and its constructs might be useful in modulating the differentiated state of cells.

## MATERIALS AND METHODS

### PC12 Cell Culture and Differentiation

PC12 cells were purchased from American Tissue Cell Culture (ATCC) and cultured in DMEM (Gibco) with 10% horse serum from PAA (Ontario, Canada), 5% FBS (Atlanta Biological, Atlanta, GA), and 1% penicillin/streptomycin. Cells were incubated in 37 °C, 5% CO_2_, and 95% humidity. Differentiation was carried out a medium of 1% horse serum, 1% penicillin/streptomycin and initiated by the addition of nerve growth factor (NGF) from Novo protein. Medium was changed every 24 h. Cells from 8 passages or less were used in these studies.

### siRNA knock-down

Down-regulation of PLCβ1 was accomplished using SmartPool rat siRNA(PLCβ1) from Dharmacon Cat# L-010280-00-0005 to give greater than 90% down-regulation as estimated by western blotting. siRNA(CDK16) from Dharmacon cat gave 86±3% down-regulation. siRNA(TRAX) was purchased from Ambion (cat# AM16708) to give 73 ± 8% knock-down, and siRNA(translin, human) from Dharmacon (cat# L-011897-01) gave 68 ± 5%. Transfection with both siRNA(TRAX) and siRNA(translin) gave (95 ± 2%). siRNA negative control sigma cat# SIC001.

### Western Blotting

Samples were placed in 6-well plates and collected in lysis buffer that included Nonidet P-40 and protease inhibitors. After SDS-PAGE electrophoresis, protein bands were transfer to nitrocellulose membrane (Bio-Rad, Inc). Primary antibodies to PLCβ1 (D-8), Gαq (E-17) were from Santa Cruz Biotechnology. Antibodies for actin and GAPDH, TRAX, CDK16 from were from Abcam, Inc. Membranes were treated with antibodies diluted 1:1000 in 0.5% BSA and washed 3 times for 5 min before applying secondary antibiotic (anti-mouse or anti-goat from Santa Cruz Biotechnology and Anti-Rabbit from Abcam at a concentration of 1:1000. Membranes were washed 3 times for 10 minutes before imaging on a Bio-Rad imager to determine the band intensities. Bands were measured at several sensitivities to ensure the intensities were in a linear range and avoid detector saturation. Data were analyzed using Image J in grayscale plot profile. Bands were normalized to loading control.

### Transfection

Plasmid transfections were accomplished using Lipofectamine 3000 (Invitrogen, Inc) as recommended by the manufacturer. Transfection of siRNAs was also carried out using Lipofectamine 3000.

### Proliferation assay

3-(4,5-dimethylthiazol-2-yl)-2,5-diphenyltetrazolium bromide (MTT; Thermo Fisher Scientific) was added to cells at a final concentration of 2mM and incubated at 37°C in 5%CO2 for 1 h. Medium was carefully aspirated from each well and replaced with 200 ml DMSO. The contents of each well were mixed by pipetting up and down and horizontally shaking the plates until the color was homogeneous. Absorbance data were obtained from PerkinElmer Victor 3 Plate Reader (Waltham, MA,USA) at 570 nm.

### miRNA sequencing

We knocked down PLCβ1 using the smartpool mixture of 4 target miRNAs from Dharmacon. The following samples were tested: Control (mock transfected), PLCβ1 knock-down, Control (mock transfected) +NGF, PLCβ1 knock-down +NGF. Each experiment had three (3) plates/sample. PC12 cells were cultured as described [9] and transfection was performed with the RNAiMAX reagent. NGF (1/1000) was added to the appropriate samples for 24 hours. Total RNA was extracted using the Qiagen miRNA Mini kit and run on a formaldehyde denaturing gel to check integrity. cDNA was synthesized for each sample using 5 µg of RNA with the Qiagen MiScript RT Kit, according to the manufacturer’s instructions. PLCβ1 knock-down was verified in each experiment by real-time PCR, using the QuantiFast SyBr Green PCR Kit (Qiagen) and the following primers: forward 5’ CGGCCAGGCTATCACTACAT 3’; reverse 3’ ATTCACCCCATTCTCTGCTG 5’. Only samples from experiments with at least 80% knock-down of PLCβ1 mRNA were used for RNA sequencing. Aliquots from the selected samples of each experimental group from three separate experiments were pooled and submitted to the University of Cincinnati Genomics Core for RNA sequencing.

### Fluorescent Oligonucleotides

A fluorescent double stranded DNA oligonucleotide based on the miR103 sequence, AGCAGCAUUGUACAGGGCUAUGA, labeled on 3’ end with 6-carbofluorescein (FAM) and the 5’ end with 5-carboxytetramethylrhodamine (TAMRA) was purchased from Genewiz. The small length of the oligonucleotide, upon excitation of FAM, Forster Resonance Energy Transfer (FRET). An oligonucleotide with 3 mismatches on the 5’ strand, AGC*GAT*AUU*T*UAC*GAA*CUAUGA also containing the same FRET pair on the 5’ and 3’ ends were also prepared.

### Confocal Imaging

Cells were seeded in poly-D-lysine coated glass-covered dishes from Mat-Tek. Images were acquired using Zeiss 510 meta confocal microscope. Data were analyzed using, LSM software and ImageJ software.

### RT-PCR

Cells were seeded in 6 well plates and transfected with siRNA of each target protein as described above. After 48 hours, cells were washed twice with PBS then lysed using Qiazol (Qiagen). Total RNA including miRNA was extracted using miRNeasy kit from Qiagen following the manufacturer’s protocol.

Reverse transcription were performed using TaqMan™ Advanced miRNA cDNA Synthesis Kit Cat# A28007 as recommended by the manufacturer. RT-PCR measurements were performed using TaqMan™ Fast Advanced Master Mix. Assays were carried out using Taqman reagents and U6 as internal control on a Roche Light cycler 96 as follows (90C for 20 Seconds then 45 cycles of 95C for 3 seconds and 60C for 30 seconds) and analyzed using Light cycler software.

### FRET/FLIM imaging

Images of cells in MatTek chambers (MatTek, Ashland, MA, USA) were acquired using a 2-photon Mai Tai laser (Spectra-Physics, Santa Clara, CA, USA) and a Nikon inverted confocal microscope in an ISS Alba System (Champaign, IL, USA). Data were analyzed using ISS Vista Vision and ImageJ (National Institutes of Health, Bethesda, MD,USA) software packages. Fluorescence lifetime imaging microscopy (FLIM) was carried out on an ISS instrument. Atto 425 (t =3.61ns) was used to calibrate the sample lifetimes.

### Microinjection studies

Cells were washed with Hanks’ Balanced Salt Solution (HBSS) media before carrying out FRET?FLIM imaging on the ISS ALBA system described above. Fluorescence lifetimes of 6-carbofluorescein (FAM) labeled oligonucleotides were excited at 780 nm. Microinjections were performed using Femtotip capillaries (Eppendorf) with 100 µM solution of the oligonucleotides injected with an injection pressure of 40 hPa over 0.65 seconds, with an Eppendorf FemtoJet 4i microinjection apparatus.

### Retraction studies

Cells were differentiated following the protocol above. Cells were stimulated by adding 5uM carbachol (Sigma) directly into the cell media while imaging.

## RESULTS

### Down-regulation of PLCβ reverses the differentiation of PC12 cells

Under typical conditions, treatment of undifferentiated PC12 cells with NGF halts proliferation and induces the growth of neurites over 36-72 hours. Here, we treated PC12 cells with NGF for 72 hours to achieve full differentiation as assessed morphologically (i.e. when the length of the neurite are greater than three times the length of the cell diameter). Cells were then washed and transfected with siRNA(PLCβ1) to reduce the level of PLCβ1 by 80-90% as determined by western blotting. In mock transfected cells, the neurite length remained constant over 72 hours but in cells treated with siRNA(PLCβ1), we find a complete retraction of neurites into the soma (**Fig. 1A-B**). Alternately, transfecting cells with siRNA(PLCβ1) 24-48 hours before NGF treatment prevented neurite growth (**Fig. 1B**) as previously noted [9].

**1.**
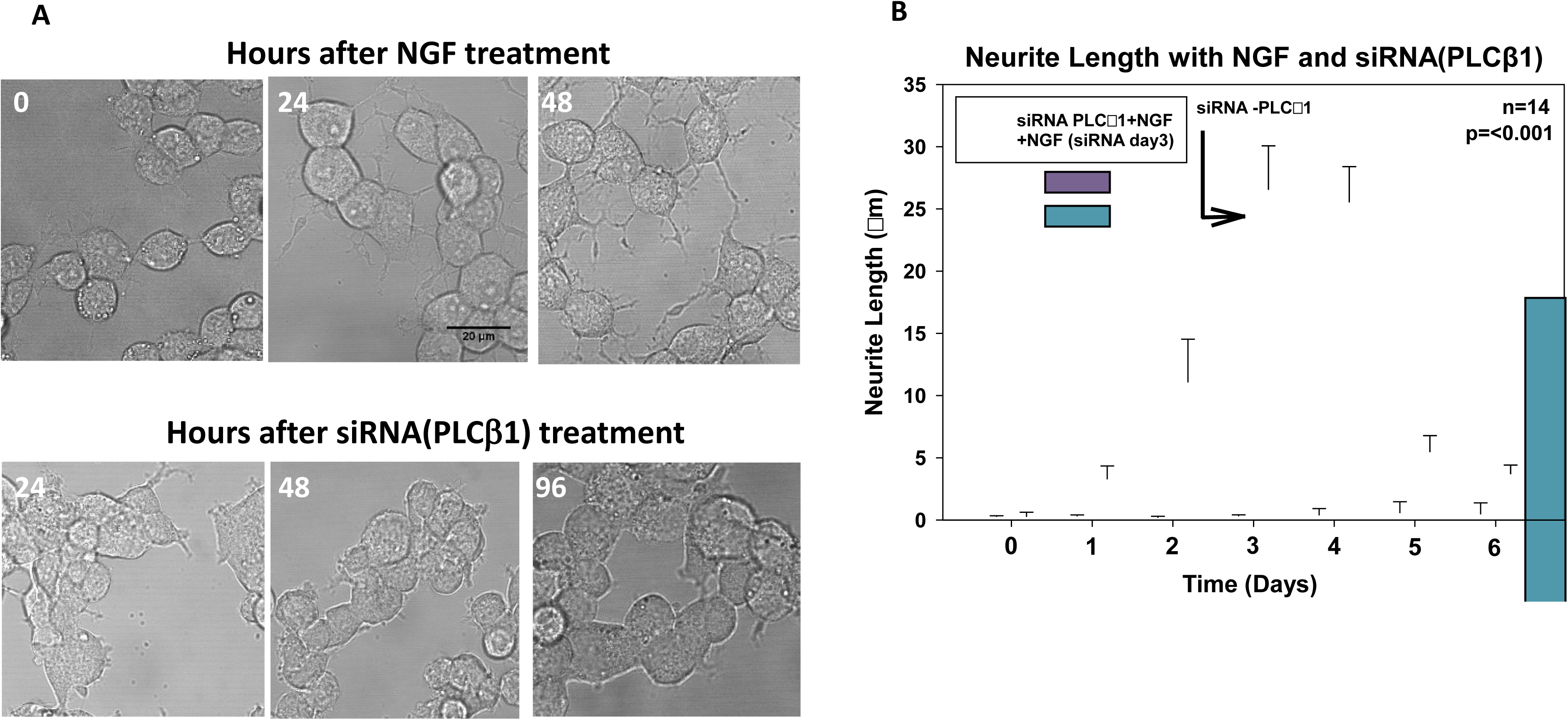
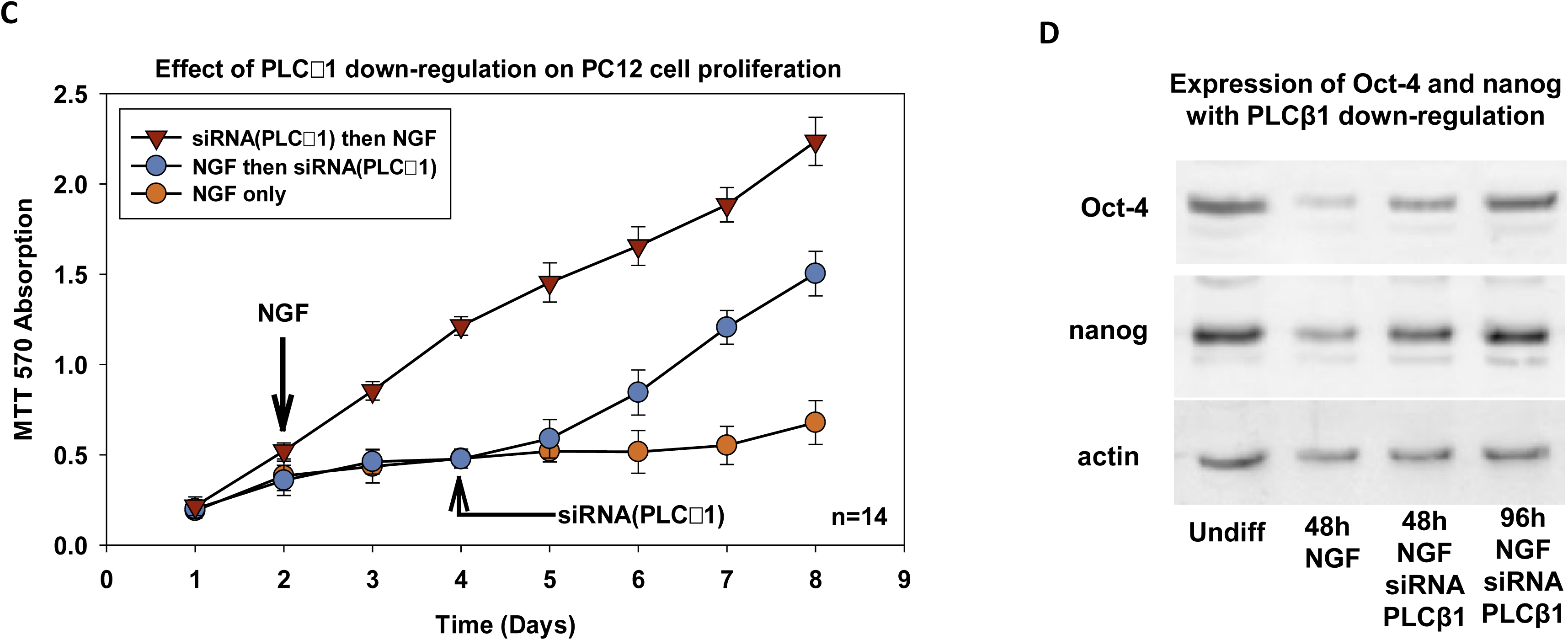
Reversal of PC12 cell differentiation with siRNA(PLCβ1). **A** - DIC images of PC12 cells (*top panels*) where the time refers to the hours after NGF treatment showing the growth of neurites. The *bottom panels* show images from the same set of cells at 48hr post NGF that were treated with siRNA(PLCβ1) showing retraction of neurites. **B –** Graph showing changes in neurite length during differentiation induced by NGF and subsequent treatment with siRNA(PLCβ1). Data are from 4 independent experiments. ANOVA comparison between 0 - 3 days and between 3-6 days is statistically significant p≤0.001. Error bars indicate SEM. **C-** MTT proliferation assay showing that NGF treated cells (orange circles) lose their ability to proliferate as they differentiate and produce neurites. However, NGF treated cells return to proliferation after siRNA(PLCβ1) (blue circles). Down-regulation of PLCβ1 before NGF treatment allows the cells to continuously proliferate (red triangles). Data are from two independent experiments and SEM is shown. **D -** Western blot showing changes in the levels of two stem cell markers, Oct4 and nanog after 48 hrs of NGF treatments, and after treating the siRNA(PLCβ1).

Differentiated PC12 cells are non-proliferative and so we determined whether down-regulating PLCβ1 would transition the cells back to a proliferative state. Thus, we monitored the proliferation of PC12 cells using an MTT assay. Down-regulating PLCβ1 before NGF treatment did not affect proliferation in contrast to cells with endogenous PLCβ1 levels (**Fig.1C**). However, transfecting non-proliferative differentiated cells with siRNA(PLCβ1) allowed the cells to resume proliferation at a rate similar to undifferentiated cells (**Fig. 1C**). Taken together, the results in **Fig.1A-C** show that PLCβ1 is required to maintain the differentiated state.

To better characterize the de-differentiated state resulting from reduced PLCβ1 levels, we monitored changes in the amounts of two stem-cell marker proteins: nanog and Oct4. In **Fig. 1D** we show that the expression levels of these proteins are high in undifferentiated cells and are greatly reduced upon NGF treatment, as expected. However, in differentiated cells where PLCβ1 is down-regulated, the levels of both nanog and Oct4 are recovered. Taken together, these studies show that loss of PLCβ1 in differentiated PC12 cells causes the cells to adapt a stem-cell like state.

### Loss of C3PO also induces PC12 cell de-differentiation

PLCβ1 has been found on the plasma membrane, the cytosol and the nucleus and we wondered which population impacts differentiation. We noted that since we are not working under conditions that activate Gαq, then it is probably not the plasma membrane population of PLCβ1, and siRNA down-regulation should effect the cytosolic PLCβ1 pool and not the nuclear one. These observations suggest that cytosolic PLCβ1 is promoting de-differentiation. As mentioned, PLCβ1 halts PC12 cell proliferation by inhibiting the mitotic kinase, CDK16, and PLCβ1 helps to induce the neurite growth by inhibiting C3PO [9, 12]. Thus, loss of PLCβ1 may promote de-differentiation due to interactions with CDK16 and/or C3PO.

We first determined whether CDK16 is involved in de-differentiation. CDK16 is primarily expressed in neuronal tissues and testis [13]. During neuronal differentiation, CDK16 transitions from its pre-mitotic role of inhibiting the proliferation inhibitor p27kip, to a post-mitotic role in synaptic vesicle trafficking (see [13]). In previous studies, we showed that PLCβ1-CDK16 associates in undifferentiated PC12 cells using Förster resonance energy transfer (FRET) [12]. However, in differentiated PC12 cells transfected with eGFP-PLCβ1 / mCherry-CDK16, we could not detect FRET (0.002±.003, n=8). Since PLCβ1-CDK16 complexes are not present in differentiated cells, it is unlikely they promote de-differentiation. In a supporting study, we de-differentiated PC12 cells expressing mCherry-CDK16 by down-regulating PLCβ1, and then transfecting with eGFP-PLCβ1. After 24 hrs we could detect both mCherry-CDK16 and eGFP-PLCβ1, but could not detect FRET suggesting that PLCβ1-CDK16 complexes do not form during de-differentiation. In a second series of studies we down-regulated CDK16 and found that treatment with NGF halted proliferation (**Fig.2A**). These results suggest that CDK16, either alone or in complex with PLCβ1, does not play a role in de-differentiation.

**2.**
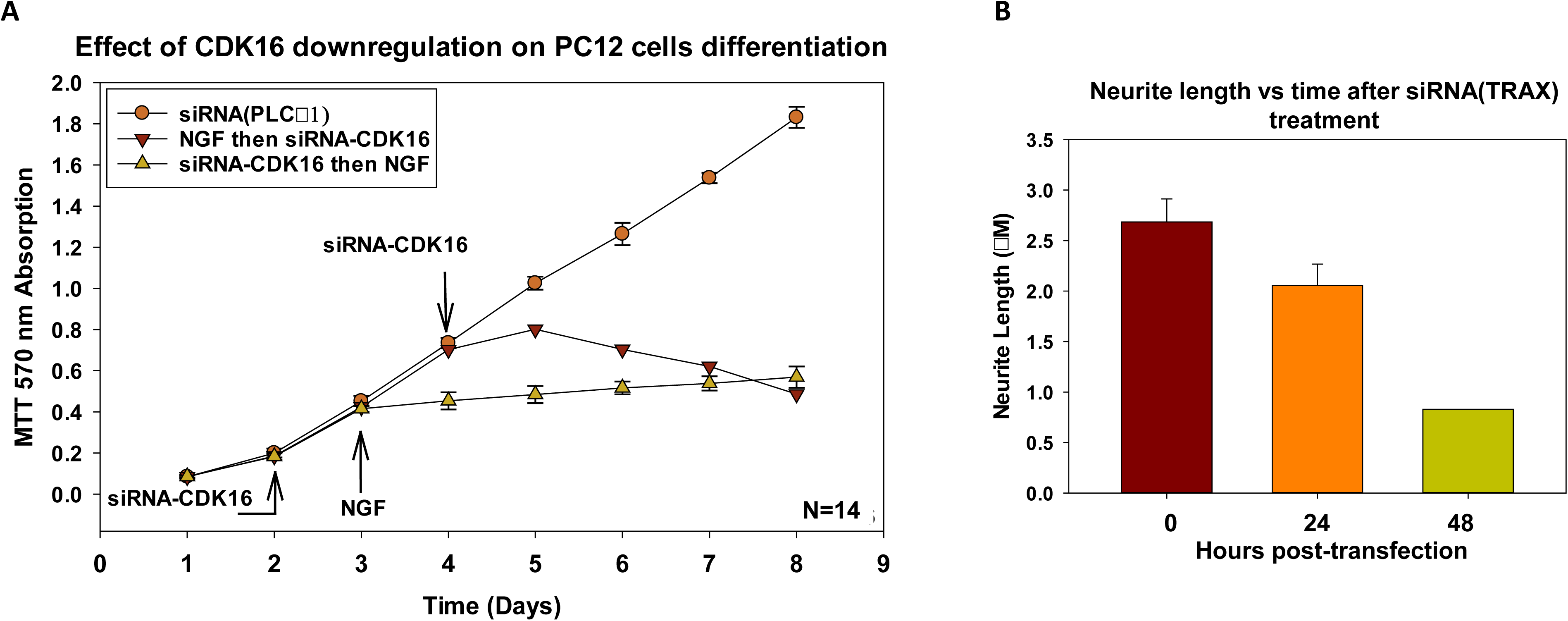
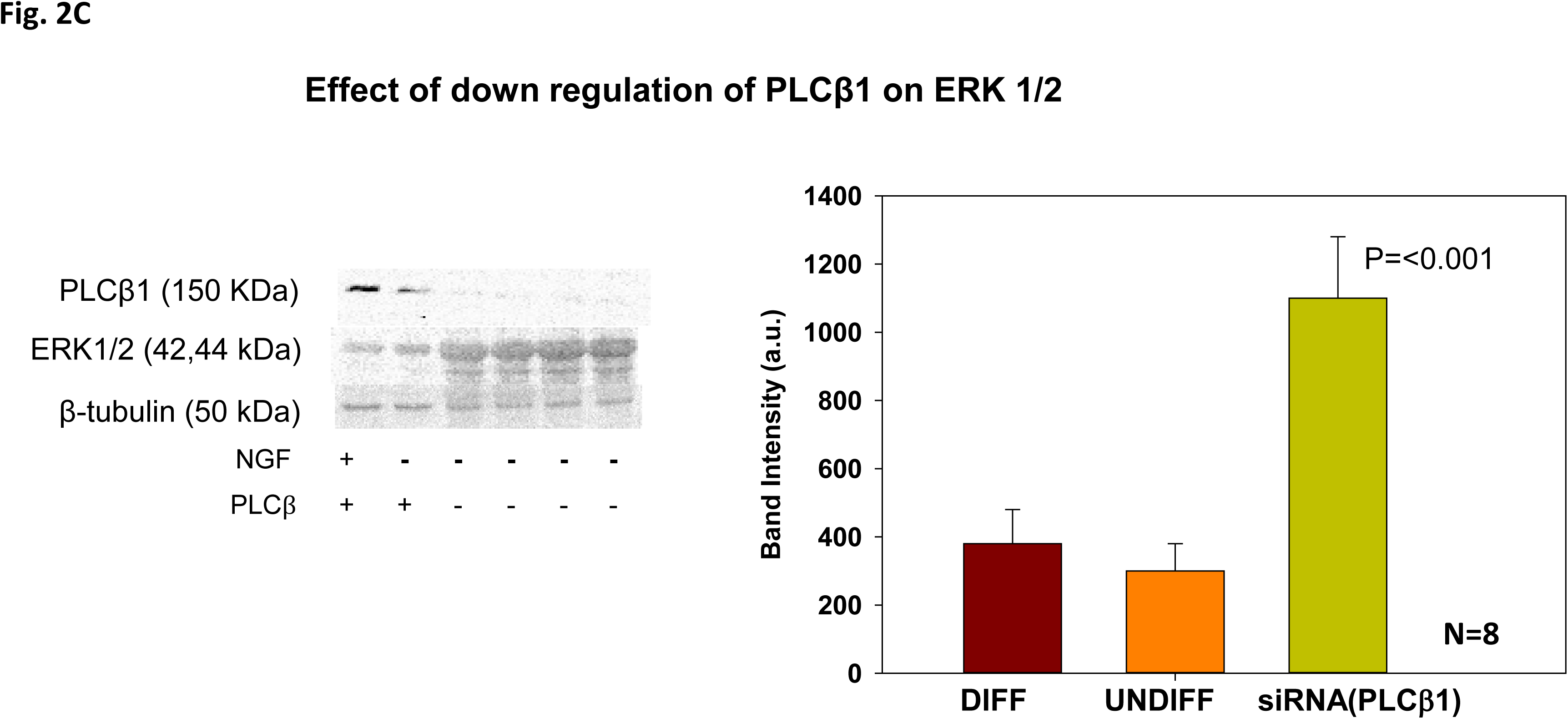
PLCβ1 down-regulation allows differentiated PC12 cells to become proliferative. **A** – Proliferation as monitored by an increase in 570nm absorption using an MTT assay. PC12 cells were treated with NGF for 48 hours to induce differentiation, and then treated with siRNA(PLCβ1) for 24 hours (orange circles), or siRNA(CDK16) (red). In an alternate study, undifferentiated cells were treated with siRNA(CDK16) on day 2 and NGF was added at day 3 (yellow triangles). These data indicate that, unlike PLCβ1, down-regulation of CDK16 does not overcome reduced proliferation by NGF treatment. **B –** The effect of TRAX down-regulation on neurite length in PC12 cells over a 2 days period where n=12. **C-** The effect of PLCβ1 down-regulation on ERK1/2 expression as determined by western blotting.

We then tested whether changes in interactions between PLCβ1 and C3PO are associated with de-differentiation. In initial studies, we down-regulated C3PO in differentiated PC12 cells. C3PO is an oligomer of two TRAX and six translin subunits, and we find that down-regulating either TRAX or translin alone results in a ∼70% reduction of C3PO. In **Fig. 2B** we show that knocking down C3PO reverses de-differentiating similar to the behavior seen for PLCβ1. These results suggest that both PLCβ and C3PO are required to maintain differentiation, and that their absence promotes de-differentiation.

If loss of PLCβ1 is associated with proliferation, we would expect to see an increase in enzymes that mediate key proliferation signaling pathways. To test this idea, we monitored the level of ERK1/2 which plays a central role in proliferative MAPK pathways [14]. We find that down-regulating PLCβ1 results in a large and significant increase in ERK1/2, supporting the direct relationship between PLCβ1 and proliferation (**Fig. 2C**).

### De-differentiation is associated with shifts in the distribution of microRNAs

C3PO has been reported to promote RNA-induced silencing [5] and its activity is inhibited by PLCβ1 [15]. With this in mind, we tested whether differentiation and subsequent de-differentiation is reflected in shifts in miR populations of PC12 cells, and whether some of these changes correlate to PLCβ1 levels. To do this, we characterized miR populations of four sets of PC12 cell samples: undifferentiated cells, undifferentiated where PLCβ1 was almost fully knocked-down (i.e >95%), differentiated cells and differentiated cells treated with siRNA(PLCβ1) for 2 days where the cells are estimated to be ∼40% de-differentiated (*supplemental*).

In comparing the miR populations of the four samples, we were surprised to find that NGF treatment did not change the total number of miRs (3.00 vs 3.04 million). However, siRNA(PLCβ1) treatment reduced the number of miRs by ∼50% in undifferentiated cells to 2.04 million and by ∼17% in de-differentiating cells to 2.57 million (*supplemental*). This loss in the number of miRs with PLCβ1 down-regulation is consistent with higher C3PO nuclease activity and greater miR processing due to the loss of inhibition of PLCβ1 (see Discussion). It is notable that the overall loss in the number of miRs with PLCβ1 is very high considering the relatively low expression of PLCβ1 in undifferentiated cells.

RNAseq studies identified 393 different miRNAs in untreated, undifferentiated cells, and of these, only 49 showed more than 10,000 copies. These are listed in **Fig. 3A**. **Fig. 3B** shows the changes in the levels of these miRs with NGF treatment. We find that the miRs whose populations change overlap with those previously identified in neuronal cell differentiation [16]. While most of these miRs showed a 10-40% reduction in the number of miRs, two of these, rno-miRs-21 and -26a, showed an increase (**Fig. 3B**). It is notable that these miRs have been associated with neuronal tissue and show a strong sequence homology (**Fig. 4A**).

**3.**
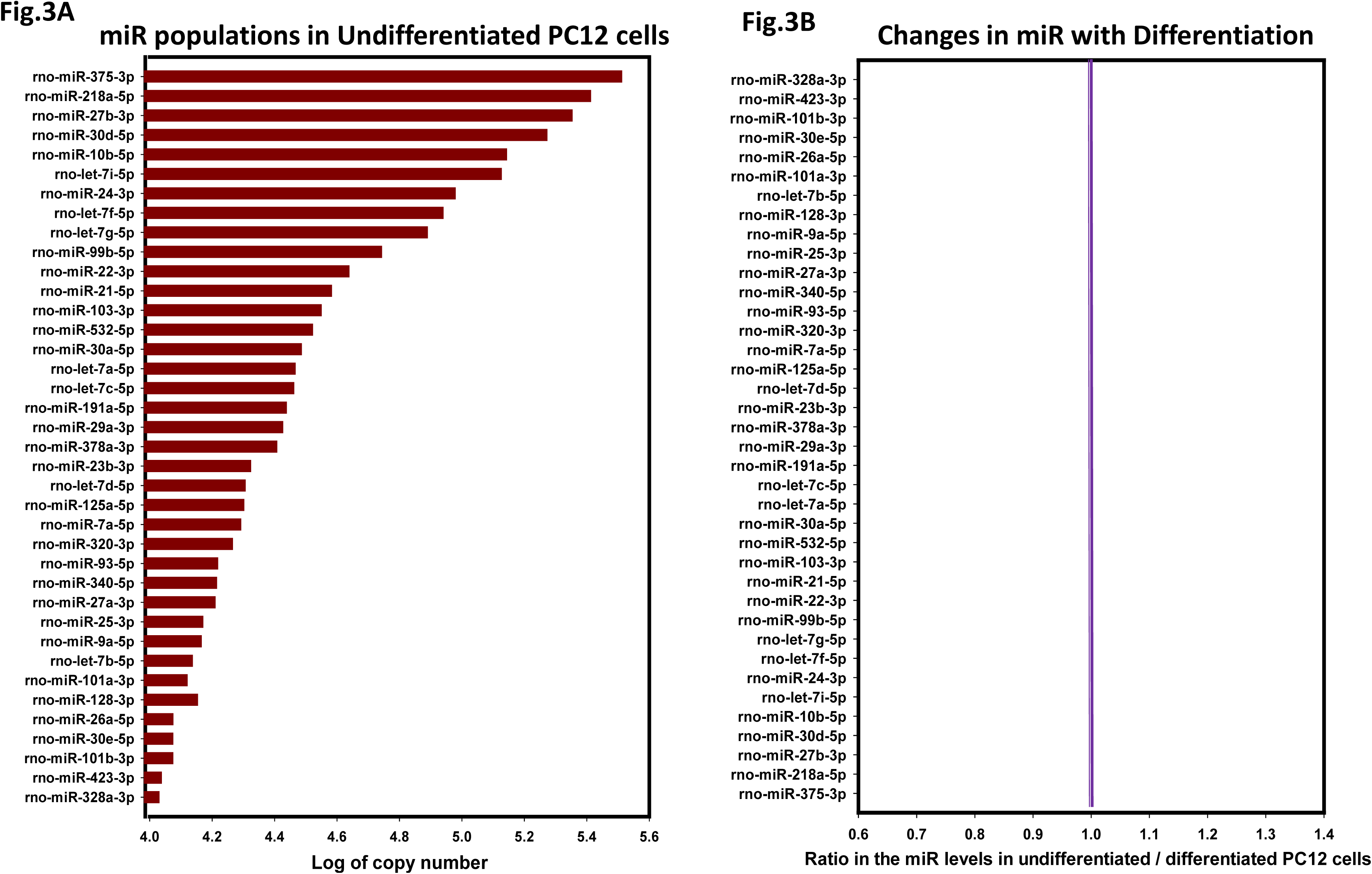
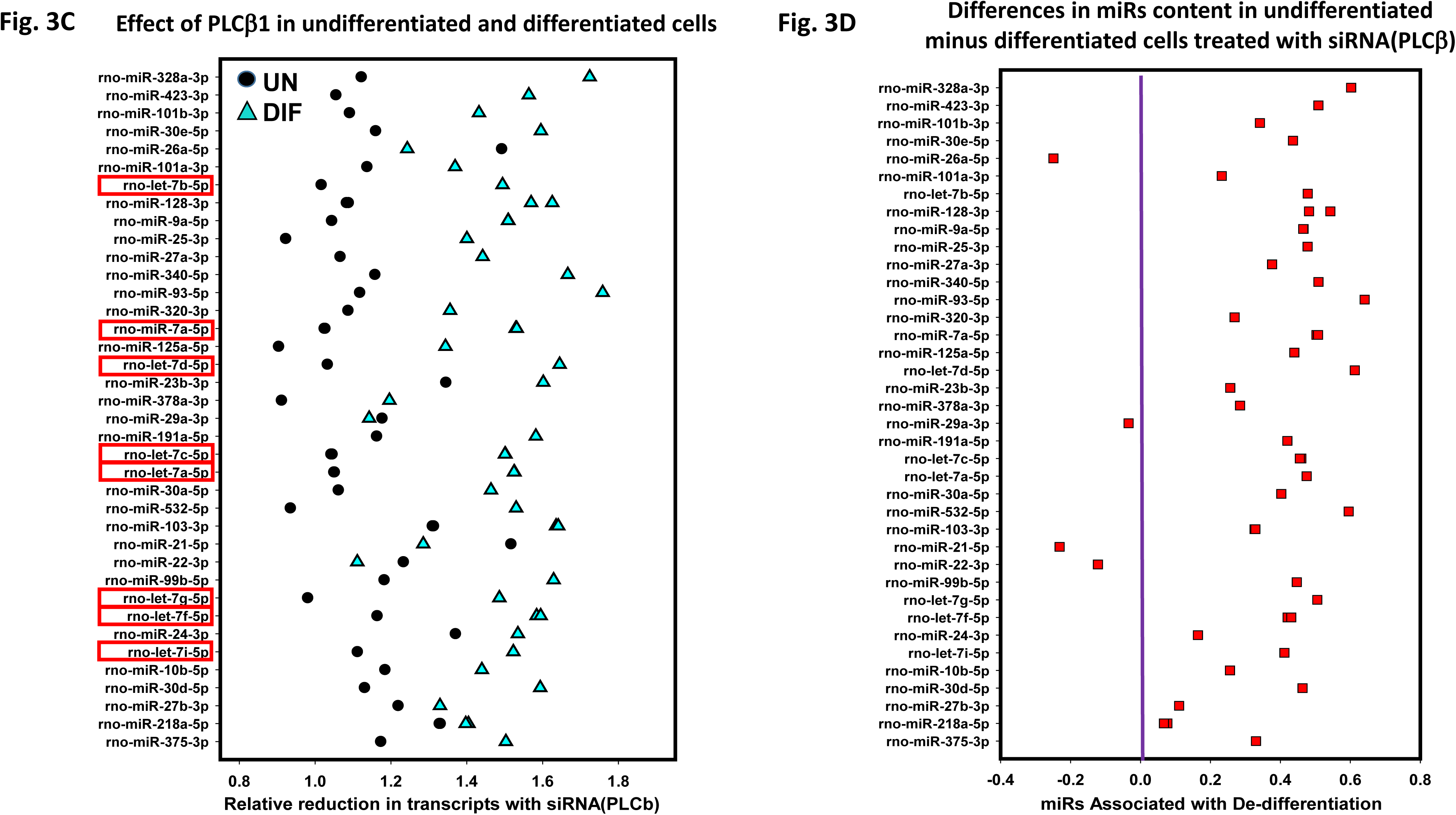
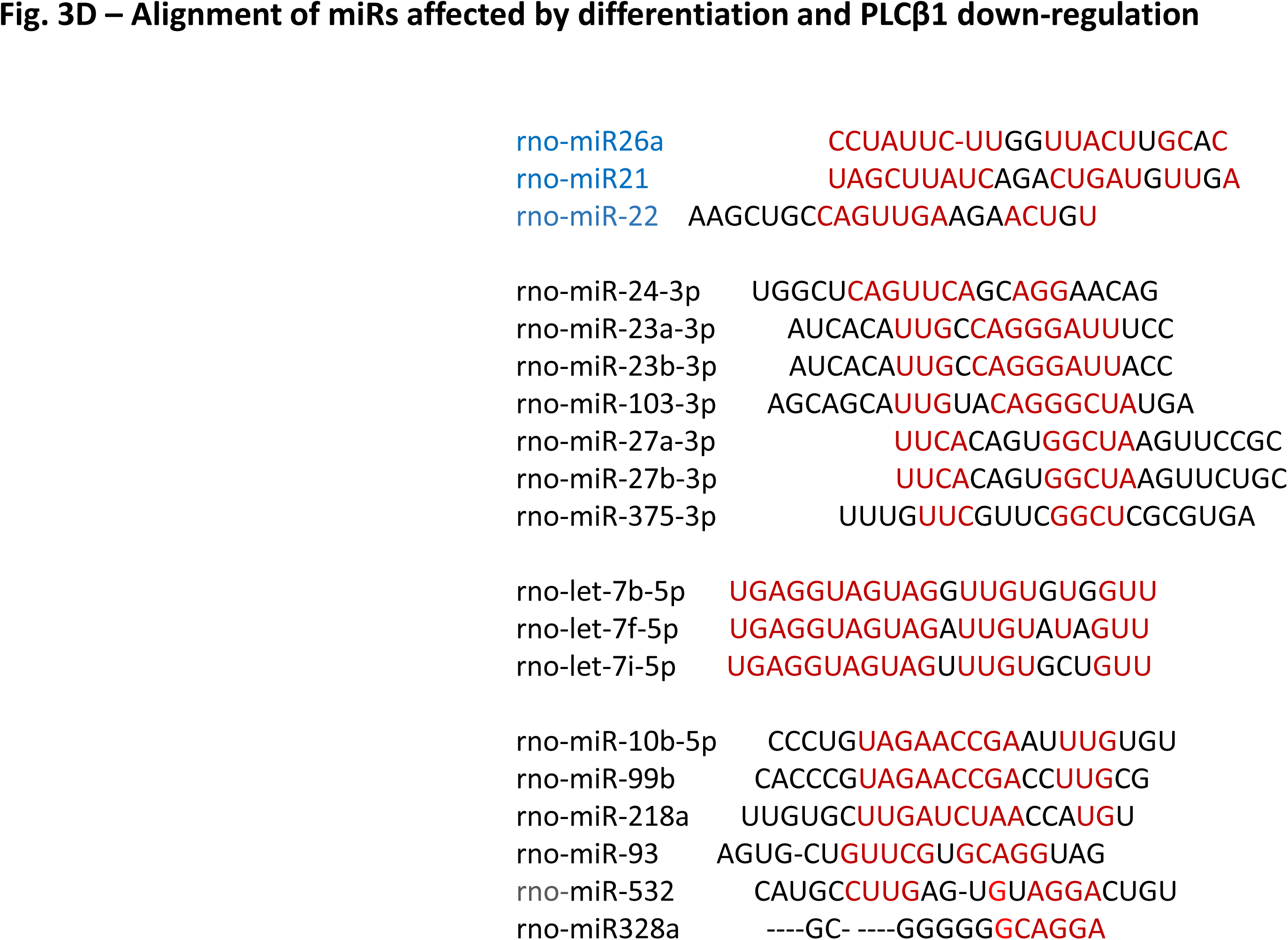
RNAseq results of PC12 cells with differentiation and PLCβ1 down-regulation. **A** – Copy number of the 49 miRs expressed in undifferentitated PC12 cells present in copy numbers greater than 10,000. **B** – Changes in the levels of the miR listed in 3A with NGF treatment as expressed as the ratio of undifferentiated / differentiated. Values below 1.0 correspnd to miRs that increase with differentiation and values above 1.0 decrease with differentiation. **C**-Changes in miR levels with siRNA(PLCβ1) where the X axis is the ratio of mock/si(PLCβ1) transfected in undifferentiated (black) and differentiated (blue). Members of the *let-7* family are identified by red boxes. **D -** To better identify the miRs involved in differentiation affected by PLCβ1, we subtracted of the undifferentiated minus differentiated values in 3C (i.e. the blue symbols minus the black). **E –** Alignment of miRs shown to be affected by PLCβ1 down-regulation. The first group (blue) are the miRs that shown an increase with differentiation. Nucleotides in red are homologous.

**4.**
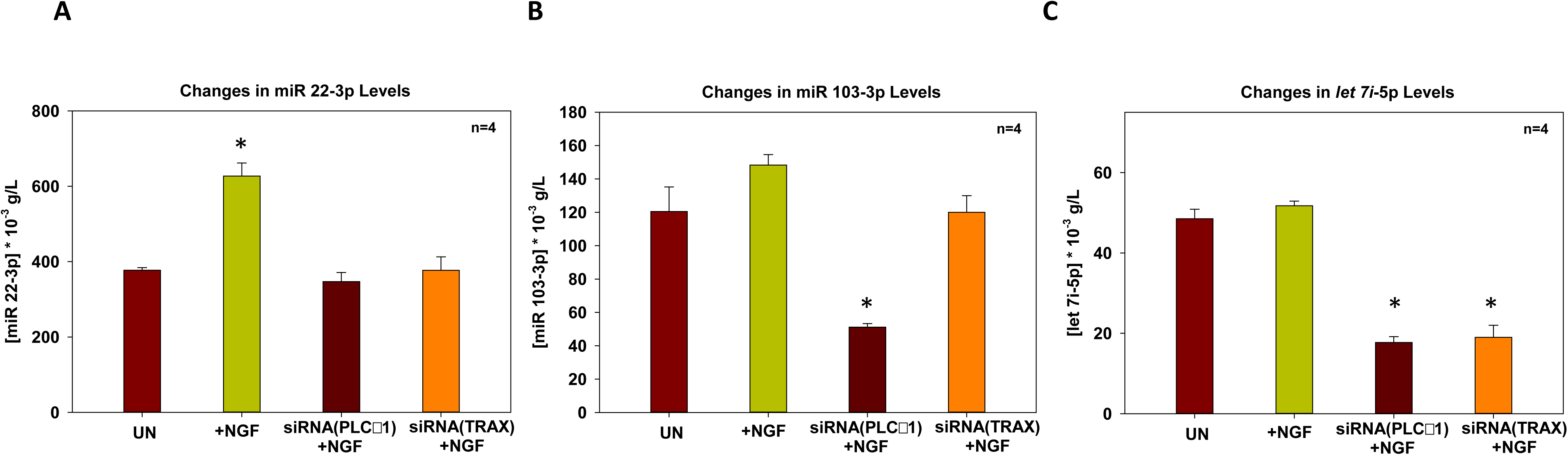
RT-PCR results of selected miRs. RT-PCR measurements were carried out as described in undifferentiated PC12 cells (UN), differentiated PC12 cells (NGF) cells and in differentiated cells treated with siRNA(PLCb1) or siRNA(TRAX) for 2 hours and where n= 4 and P≤0.001 and SD is shown.

In **Fig. 3C**, we compare the miRs that are impacted when PLCβ1 is down-regulated in undifferentiated and de-differentiated cells. In undifferentiated cells, loss of PLCβ1 decreases the number of individual miRs between −10 and 30% (black circles) consistent with increased C3PO activity. In de-differentiating cells, down-regulating the enzyme decreases the levels of almost all miRs between 5-50% (blue triangles). To more easily compare these data, we subtracted the transcripts reduced by PLCβ1 down-regulation in de-differentiating cells from undifferentiated cells (**Fig. 3D**) (i.e. the blue minus the black points in **3C**). While we see that almost all the transcripts increase during the de-differentiation process, miRs-21 and miR-26a, are upregulated and miRs-22 and -29a to a lesser extent (**Fig. 3D**).

The data in **Fig. 3** only focus on the miRs with high copy numbers. Many miRs with lower copy numbers change with differentiation and PLCβ1 knock-down. A few miRs show higher than a two-fold change with differentiation and/or PLCβ1 knock-down but their copy numbers are very low (i.e. <50) which may accentuate changes that are not significant, and so these miRs are not considered. In a separate analysis, we screened the data for families of miRs that change at least 25% and are present at 200 copy number or higher (counting the -5p and -3p as a single type) (*<u>supplemental</u>)*. By analyzing this expanded set of data, we find that 17 members of the let-7 family are reduced with PLCβ1 down-regulation, and the eight with the highest copy number are shown in red boxes in **Fig. 3C**. Additionally, the miR-30 family has 10 members that change, miR-125 has 5 members, miR-344 has 5 members, and miR-99, miR-181 and miR-130 families all have 4 members that change with PLCβ1 down-regulation.

We aligned the mature miRs of these different families to determine common sequences. These families include one group of up-regulated miRs, the let-7 family, and two other down-regulated groups. While there was no overall homology, if we separate these into different groups, some homologous groups emerge (**Fig. 3D**). It is notable that none of these groups are significantly different in terms of physical properties, such as Tm or GC content, and none fit into specific miR clusters. While some sequence homology in the groups are seen, in general, this analysis indicates a lack of strong sequence specificity in miRs and their changes in levels with de-differentiation.

We used RT-PCR to verify the results for a few representative miRs (**Fig. 4**). We find that in control cells, miR-22-3p increases ∼1.8 fold when treated with NGF where the number of transcripts significantly increase with NGF treatment, and these levels return to basal with PLCβ1 or C3PO down-regulation. These results closely follow the RNAseq results. We also tested a representative miR that does not change greatly with differentiation, let-7i-5p, and find that these levels show a small increase with differentiation, and a reduction with PLCβ and TRAX knock-down, again in accord with the RNAseq. In addition, miR-103 shows a small increase with differentiation but a significant decrease with both PLCβ1 and C3PO down-regulation. Taken together, these results corroborate the RNAseq studies that show only minor changes in miR levels with differentiation except for a few miRs, such as miR-22, and down-regulation of PLCβ1 greatly reduces the number of miRs. Additionally, our studies show that C3PO down-regulation produces the same changes as PLCβ1 supporting the idea that PLCβ1 works with C3PO in regulating miR populations to control the differentiated state.

### Cross-regulation of PLCβ by *let-7*

Because PLCβ1 levels are inversely correlated to profileration, but positively correlated to the *let7* levels, which in turn represses proliferation, we tested the possibility of reciprocal regulation of PLCβ1 by members of the *let-7* family. This study was carried out by transfecting cells with *let-7a* and monitoring the relative changes in the levels of PLCβ1. We simultaneously monitored H2B levels to follow proliferation. We find that treating cells with siRNA(PLCβ1) for 2 days increases the amount of H2B consistent with increased proliferation (**Fig.5**). However, transfecting control cells with *let-7a* reduces PLCβ1 levels and this reduction no longer occurs when C3PO is down-regulated regulation. Taken together, these results suggest an interdependence between PLCβ1, C3PO and *let-7a* in cell proliferation.

**5.**
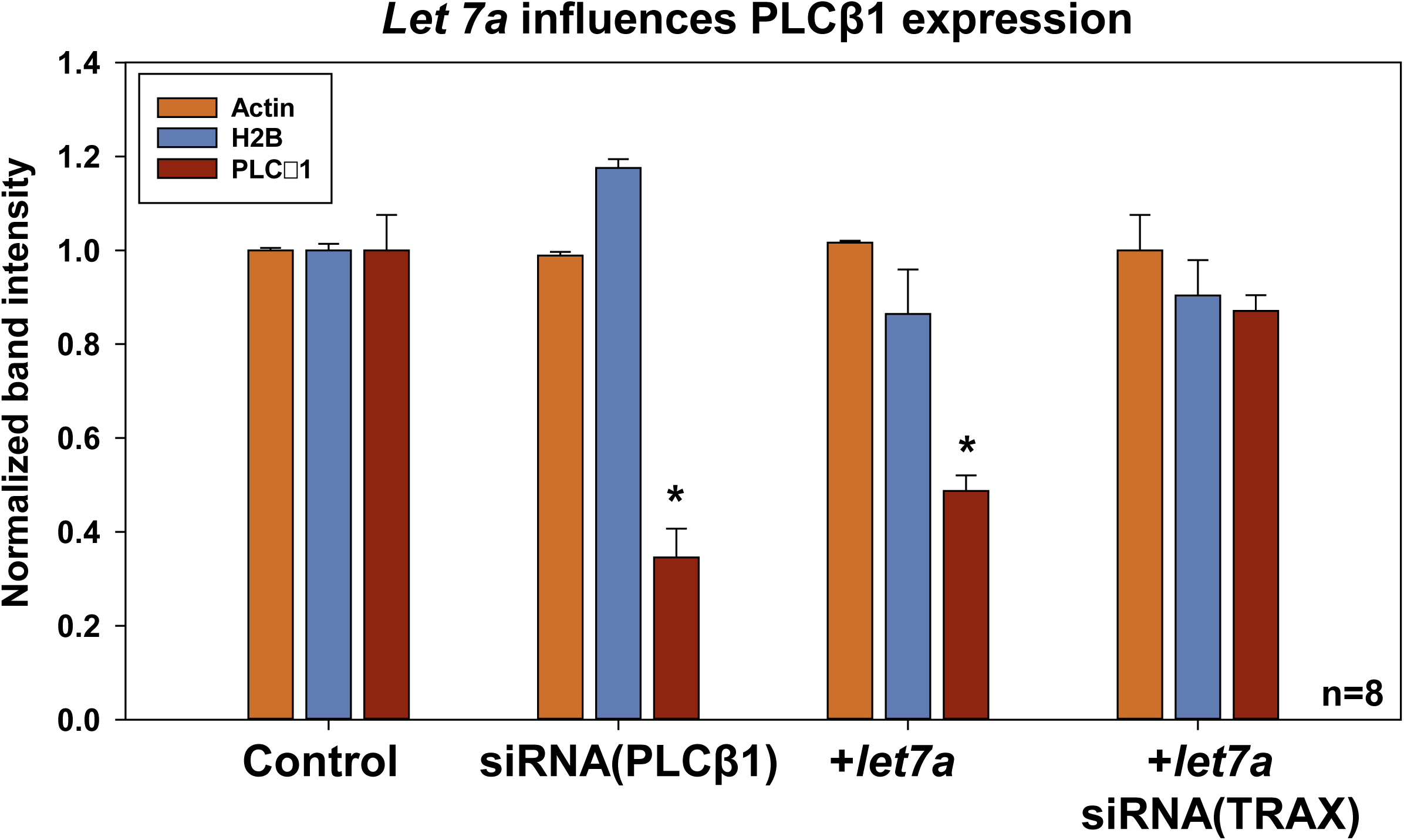
*Let-7a* influences PLCβ1 expression. Compilation of band intensities from western blots showing that transfection of *let-7a* in undifferentiated PC12 cells reduces endogenous levels of PLCβ1 and down-regulation of TRAX restores PLCβ1 levels, where n=8, * P≤0.001 and SD is shown.

### Neurotransmitter stimulation can promote the return of differentiated PC12 cells to a stem-like state

We determined whether stimulation of Gαq could shift the cytosolic population of PLCβ1 to the plasma membrane and away from C3PO to reverse differentiation. Stimulation of Gαq results in an increase in intracellular calcium within the first minute followed by recovery over the next hour. Concurrent with this recovery is internalization of muscarinic receptors allowing for desensitization [17].

Starting with fully differentiated PC12 cells, we followed changes in morphology over 30 minutes with under continuous exposure to carbachol (a derivative of acetylcholine). Immediately upon stimulation, we see retraction of neurites towards the soma membrane resulting in full retraction after ∼10 minutes resulting in cells with an undifferentiated-like morphology (**Fig 6A**). To better understand the properties of these retracted cells, we measured changes in the levels of nanog in cells stimulated with carbachol for 30 minutes. We find a slight recovery in nanog levels consistent with cells that are transitioning out of the differentiated state (**Fig. 6B**). In a corroborative study, we measured changes in two different miRs (miR-22 and 23a) with differentiation and under carbachol stimulation (**Fig. 6C**). In line with the RNAseq studies, the levels of both miRs both change with differentiation, and carbachol stimulation causes their levels to shift towards those seen in undifferentiated cells.

**6.**
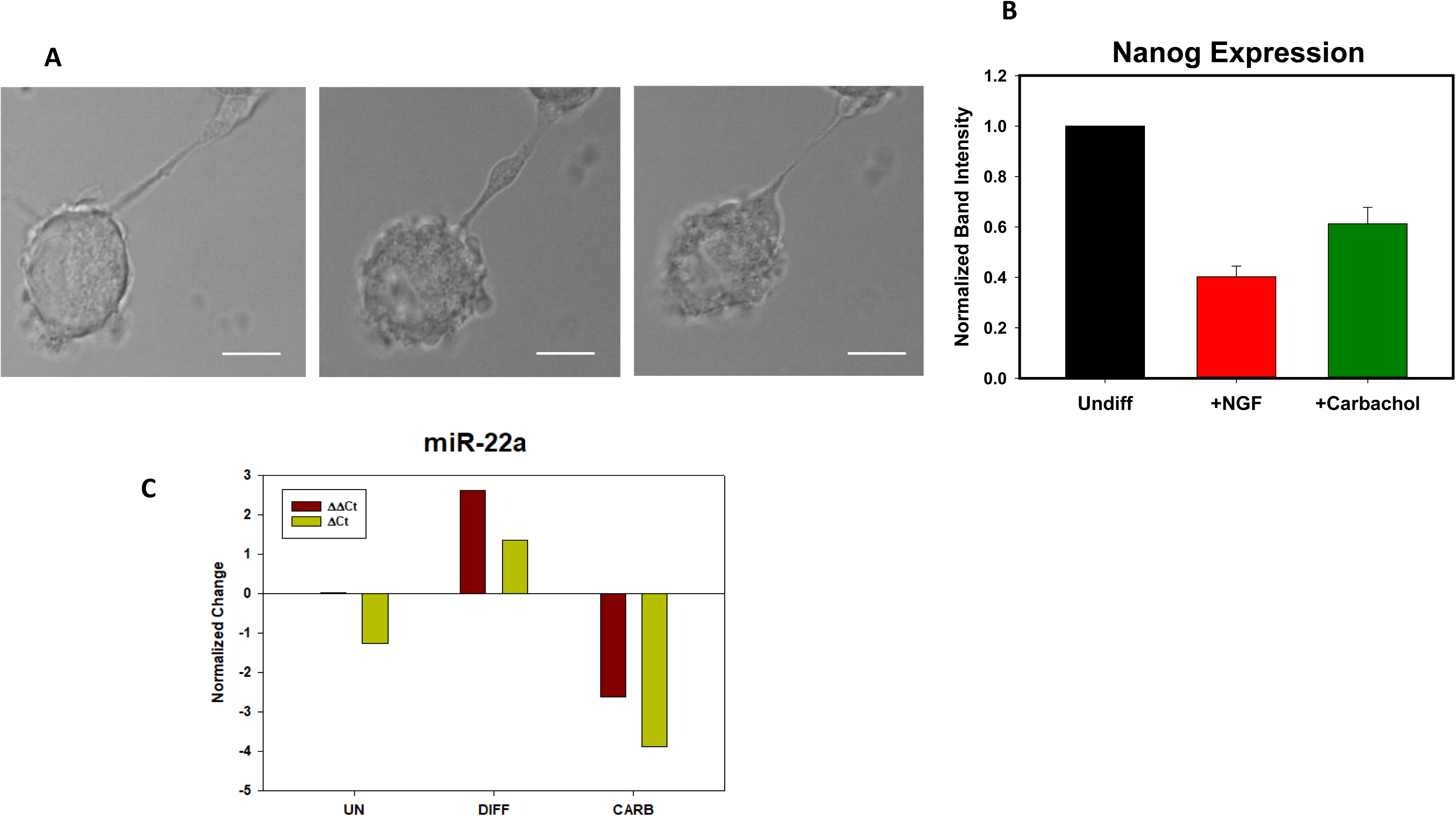
Gαq stimulation causes neurite retraction. **A –** Images of differentiated cells that have been transfected with mCherry actin, at time 0, time ∼10 minutes and ∼30 minutes after stimulation with 5μM carbachol. Scale bar is 20μM. **B-** Normalized band intensities from western blots of differentiated PC12 cells showing changes in nanog expression 30 min after carbachol (5μM) where n=2 and SD are shown. **C-** Changes in ΔΔCt values for miR-22 (*left*) and miR-23a (*right*) of differentiated PC12 cells (DIFF) under basal conditions and 30 min 5 µM carbachol stimulation (CARB) normalized to undifferentiated controls (UN).

### Live cell imaging of miR processing

Thus far, our measurements have assessed changes in miR levels in cell lysates and we wanted to assess miR processing in intact, living cells. Our approach was to monitor the degradation of miR-103 labeled on the 5’ and 3’ ends with a FRET donor / acceptor pair (FAM/TAMRA) and introduce them into cells by microinjection. This oligonucleotide has a high degree of FRET that decreases as it degrades with nuclease activity. In these measurements, we monitored FRET by the decrease in fluorescence lifetime of the donor (FAM) due to energy transfer to acceptor (TAMRA). Hydrolysis of the oligonucleotide is seen by a net increase in fluorescence lifetime. For comparison, we also constructed an imperfect miR103 analog with 3 mismatched nucleotides. This imperfect nucleotide should result in stalled Ago2 complexes[18] and should not be sensitive to changes in the levels of PLCβ1 or C3PO.

In **Fig. 7** we show the shift in the fluorescence lifetime of miR103-3p labeled with the FAM/TAMRA FRET pair at 3 minutes in differentiated cells and note that similar results are observed in undifferentiated cells. Additionally, the values remain constant over the 3-20 minutes viewed. Cell images show that the oligonucleotide is not distributed uniformly in the cytosol but appears to sequester in aggregates, which may correspond to RISC. Microinjecting the tagged miR into PC12 cells showed a clear increase in lifetimes from 1.4 to 2.8 ± 0.13 ns in undifferentiated cells and 1.8 to 3.2 ± 0.097 ns in differentiated cells showing that this miR is processed slightly more efficiently in differentiated cells. When we down-regulate PLCβ1, we find a pronounced shift to longer lifetimes, consistent with increased C3PO activity. In contrast, when we down-regulate TRAX (**Fig. 7**), hydrolysis is reduced. Processing was greatly reduced with carbachol stimulation, which is consistent with reduced levels of cytosolic PLCβ1 and sequestration of the miRs in stress granules (*see discussion*). In contrast, the imperfect miR tended to sequester in the nucleus, and both the cytosolic and nuclear populations were not impacted by changes in the levels of PLCβ1, C3PO or carbachol.

**7.**
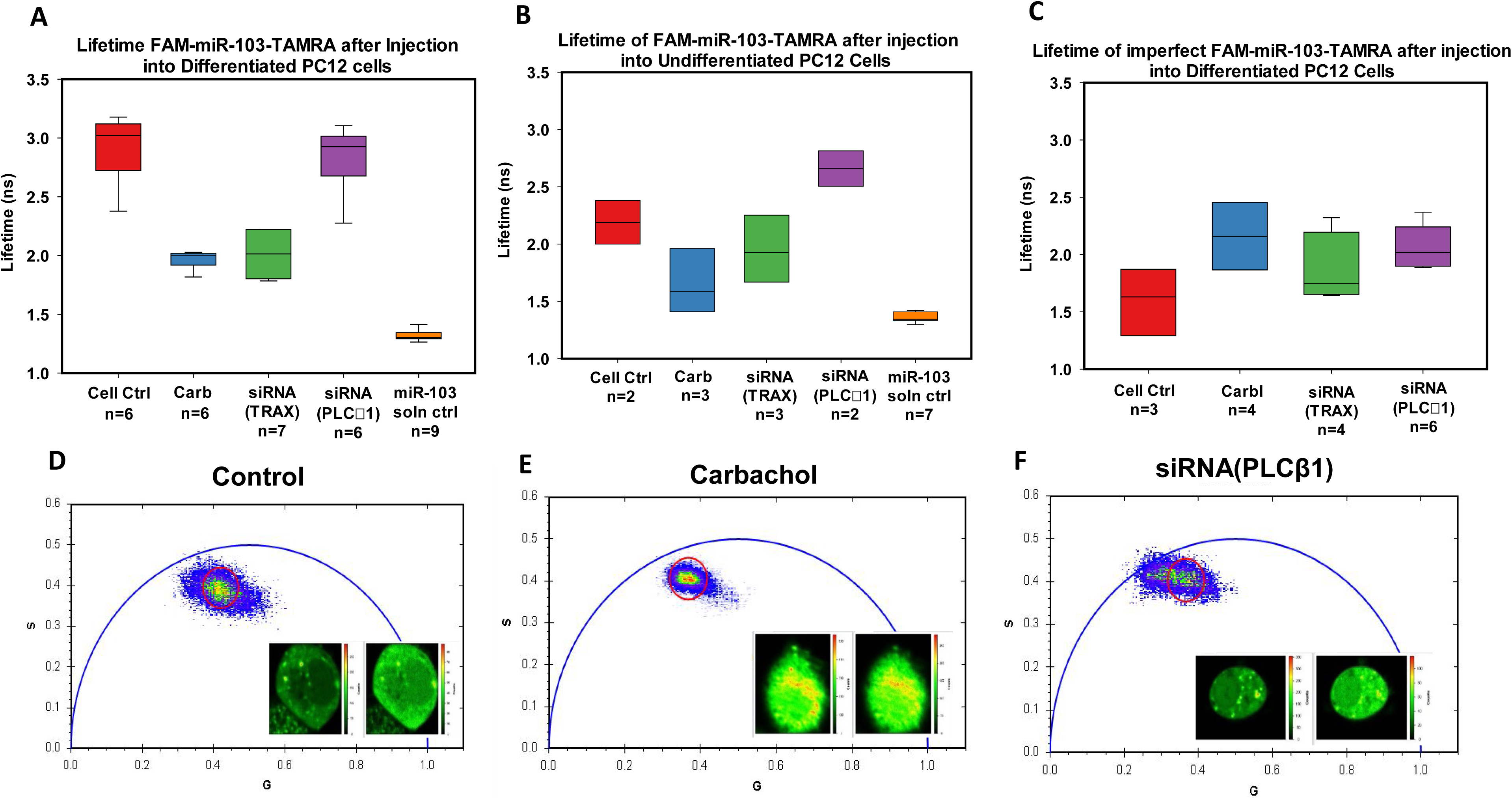
FLIM measurements in live cells show miR processing. Studies showing live processing of miR-103-3p labeled on the 3’ and 5’ ends with the FRET pair FAM/TAMRA. Here, hydrolysis of the oligonucleotide is followed by the increase in FAM lifetime from dequenching by TAMRA. In all graph, microinjection into untransfected cells is labeld *Ctrl*, stimulation with 2 μM carbachol after microinjection is labeled as *Carb*. Graphs representing lifetimes of fluorescently labeled miR-103 in differentiated **(A)** and (**B)** undifferentiated PC12 cells. **C** –Identical study as in (**A**) but using an imperfectly paired version of FAM-miR-103-TAMRA. **D –** Example of a phasor plot and the corresponding image from a FLIM measurement of FAM-miR-103-TAMRA expressed in wild type undifferentiated PC12 cells. The heat map indicates FAM and TAMARA signal intensity. **E** – Similar study as (**D**) but with undifferentiated cells stimulated with 2 μM carbachol after injection. **F** – Similar study as (**D**) but with undifferentiated cells transfected with PLCβ1 siRNA before injection.

## DISCUSSION

In this study, we show that we can control the differentiation of a model neuronal cell line by changing the levels of cytosolic PLCβ1 either by varying its expression or by driving it to the membrane with stimulation of Gαq. Specifically, our studies suggest that while PLCβ1-C3PO complexes are required to promote the transition of PC12 cells to the differentiated state, a basal amount is required to maintain differentiation. Our studies show that PLCβ1 levels regulate miRs associated with differentiation, and this regulation is in part associated with PLCβ1’s ability to inhibit C3PO.

A yeast two-hybrid study found that C3PO is a binding partner of PLCβ1 and this association was verified by coIP, pull-downs, co-immunofluorescence and FRET in all cell lines tested (HEK293, SK-N-SH, HeLa and PC12 cells [7, 8]). C3PO has been reported to promote RNA-induced silencing by processing the passenger strand of silencing RNA after an initial nick by Ago2 which leaves the guide strand free to hybridize to its target mRNA and be degraded by Ago2 of RISC [5]. Because the nuclease activity of Ago2 is operative when the miR-mRNA is perfectly paired, C3PO will only promote silencing in cases of perfect pairing between miR and mRNA, or in exogenous siRNAs. Support for this model comes from our live cell imaging data showing that hydrolysis of perfectly paired miR103-3 depends on the level of C3PO, PLCβ1 but imperfectly paired miR103-3 does not. It is notable that that PLCβ1 inhibits C3PO processing of miRs with higher Tms that are more slowly hydrolyzed by C3PO [15]. Thus, the impact of PLCβ1 on RNA-induced silencing by C3PO inhibition is limited to specific endogenous miRs.

In previous work, we followed the differentiation of PC12 cells stimulated by the neurotropic agent NGF, and found that both PLCβ1 and C3PO are required for neurite growth. Undifferentiated PC12 cells, like most stem cells, have a very low expression of Gαq and PLCβ1 since these cells have not developed mature response networks to neurotransmitters. Within the first 24 hours after NGF treatment, the level of PLCβ1 increases while Gαq and C3PO remain constant. Our studies show that this increased PLCβ1 associates with C3PO. This period is then followed by increased Gαq production and a loss of PLCβ1-C3PO complexes. However, some PLCβ1-C3PO complexes remain in the differentiated state. Although other mechanisms are possible, we postulate these complexes help maintain the differentiated state by modulating miR populations.

We propose that the ability of PLCβ1 to regulate differentiation can be traced to its regulation of C3PO, and in turn, modulation of silencing by miRs. Therefore, we carried out RNAseq in undifferentiated and differentiated cells, and in undifferentiated cells where PLCβ1 was almost completely absent, in cells undergoing de-differentiation through down-regulation of PLCβ1. While the total amount of cellular miR was unchanged with differentiation, there were some shifts in the populations of several species (see **Fig. 3B**), and these are in line with those reported in other studies of developing rat and monkey brains [16]. Notably, two species showed a substantial increase with differentiation, miR-21 and mi-26a. In contrast to our data, miR-21 levels are positively correlated with proliferation in cardiac smooth muscle tissue [19], while miR-26a is inversely correlated with proliferation in a number of tissues correlation with our results (for review see [20]). The levels of many other miRs were found to decrease slightly with differentiation. Of note is the relatively large decrease in miR-93 whose overexpression diminishes PTEN activity and promotes carcinogenesis through dysregulation of the PI3K/AKT pathway [21].

In general, our data show that reducing PLCβ1 substantially decreases the total amount of cellular miRs. Reduction of total cellular miRs with PLCβ1 down-regulation is in line with the large increase in small RNAs seen when PLCβ1 is overexpressed in HEK293 cells [7]. While this observation is consistent with increased C3PO activity, it is probable that other downstream mechanisms contribute considering the limited set of miRs that are expected to be impacted by PLCβ1-C3PO association. Of note is the strong dependence of the *let-7* family with PLCβ1 expression is in accord with the impact of *let-7* on signaling pathways (e.g. [22]). We find that the levels of 17 members of the *let-7* family changed with PLCβ1 expression. In nematodes and drosophila, *let-7* is a key regulator in differentiation and development, and in vertebrates, *let-7* levels are associated with differentiated state [23] which correlates well their involvement in de-differentiation. The interdependence between PLCβ1 and *let-7* implies a feedback mechanism that asserts multiple controls on proliferation and provides a roadmap into potential mechanisms that might drive differentiation processes initiated by changes in PLCβ1 expression. It is notable that C3PO down-regulation does not impact proliferation, as seen by H2B levels.

In addition to *let-7*, several other miR families are associated with PLCβ1-induced de-differentiation. We searched for commonalities between the miRs that increase or decrease with PLCβ1 down-regulation, we could not find a strikingly strong correlation even when divided into groups. However, we note that the local copy number of these miRs and the presence of competitive ones may override absolute sequence dependence making it unclear whether common regions found in the alignment of these groups these are true targets of C3PO-PLCβ1. Broadly, we propose that the global shift in the population of these miR families that not only affect the translation of specific mRNAs, but also function to compete with other miR to indirect affect protein production. Future systems biology studies that probe the mRNAs and proteins regulated by these miRs will offer better insight.

Our studies show a surprising relationship between activation of Gαq and de-differentation through PLCβ1. Upon Gαq activation by carbachol, we find that the neurites of differentiated PC12 cells rapidly retract resulting in a morphology that resembles undifferentiated cells. Our studies indicate that the levels of miRs begin to return to undifferentiated ones although the timeline appears to be much longer. Our studies also show that carbachol reduces the hydrolysis of the fluorescent-tagged miR103 and this observation correlates well with studies showing that reduction of carbachol stimulation increases the number and size of stress granule particles [24]. The link between Gαq and stress granules is through the ability of cytosolic PLCβ1 to bind stress granule proteins inhibiting their aggregation. We propose that the formation of stress granules caused by Gαq activation incorporates miR103 into the aggregates thereby protecting the nucleotide from hydrolysis. It is likely that a similar mechanism occurs for other miRs impacted by PLCβ1. Taken together, our results show a clear connection between external agents that stimulate and differentiation.

Our studies used PC12 cells as a model for differentiation and since the Gαq/PLCβ pathway is ubiquitous in mammalian cells, and PLCβ1 and C3PO have been found to associated in other cell lines then it is possible that link between PLCβ1 and stemness, and feedback from the extracellular environment can control the level of differentiation is a general. These finding suggest that PLCβ1 constructs could be used a method to control the degree of the level of differentiation.

## ACKNOWLEDGEMENTS

The authors would like to thank Dr. V. Siddartha Yerramilli for his helpful comments throughout this work and to Dr. David McKinnon for help with the RNAseq studies. This work was supported by NIH GM116187. OG was supported by funds from the Richard Whitcomb award.

## Supplementary Data

**Osama Garwain, Barbara Rosati, Nathan Johnson, Dmitry Korkin and Suzanne Scarlata**
| #miRNA | DM14.1 (Control) | DM14.2(Control + NGF) | DM14.3(PLCb1 KO) | DM14.4(PLCb1 KO + NGF) | 1vs3(%) | 2vs4(%) | 1v3(log2) | 2v4(log2) |
| --- | --- | --- | --- | --- | --- | --- | --- | --- |
| rno-miR-375-3p | 312450 | 305234 | 207853 | 260195 | 0.40206188 | 0.15930913 | 0.58806177 | 0.23032243 |
| rno-miR-218a-5p | 248358 | 266035 | 176830 | 200345 | 0.33645352 | 0.28170162 | 0.49005816 | 0.40912956 |
| rno-miR-218a-5p | 239358 | 257236 | 171309 | 193346 | 0.3314072 | 0.28358878 | 0.48256908 | 0.41190764 |
| rno-miR-27b-3p | 216880 | 237451 | 163146 | 194758 | 0.28279118 | 0.1975572 | 0.4107334 | 0.28594724 |
| rno-miR-30d-5p | 180048 | 160389 | 112969 | 141869 | 0.45785057 | 0.12254432 | 0.67245464 | 0.17701582 |
| rno-miR-10b-5p | 133718 | 149239 | 92864 | 126030 | 0.36061117 | 0.16862778 | 0.52600235 | 0.24385741 |
| rno-let-7i-5p | 128640 | 122557 | 84470 | 110262 | 0.41452771 | 0.10561853 | 0.60682836 | 0.15251721 |
| rno-miR-24-3p | 91543 | 94548 | 59641 | 68990 | 0.42202879 | 0.31256344 | 0.61814512 | 0.45465968 |
| rno-miR-24-3p | 91536 | 94543 | 59640 | 68986 | 0.42197174 | 0.31256841 | 0.61805899 | 0.45466703 |
| rno-let-7f-5p | 83682 | 89941 | 52824 | 77301 | 0.452112 | 0.1511582 | 0.66372378 | 0.21849185 |
| rno-let-7f-5p | 82726 | 88626 | 51871 | 76158 | 0.45847976 | 0.15132537 | 0.67341264 | 0.21873441 |
| rno-let-7g-5p | 74580 | 66680 | 50190 | 68009 | 0.39095937 | -0.0197344 | 0.57138885 | -0.0284716 |
| rno-miR-99b-5p | 53187 | 56734 | 32649 | 47991 | 0.47854047 | 0.16697064 | 0.70403486 | 0.24144971 |
| rno-miR-22-3p | 41795 | 47486 | 37611 | 38508 | 0.10538246 | 0.20880527 | 0.1521757 | 0.30234404 |
| rno-miR-21-5p | 36746 | 56315 | 28590 | 37129 | 0.24966328 | 0.41064167 | 0.36207659 | 0.60097279 |
| rno-miR-103-3p | 34065 | 35400 | 20827 | 27032 | 0.48232894 | 0.26806766 | 0.70983516 | 0.3890811 |
| rno-miR-103-3p | 33773 | 35046 | 20578 | 26699 | 0.48554764 | 0.27037007 | 0.71476757 | 0.39246408 |
| rno-miR-532-5p | 31957 | 27067 | 20884 | 28951 | 0.41910638 | -0.0672641 | 0.61373392 | -0.0970782 |
| rno-miR-30a-5p | 29428 | 27481 | 20103 | 25892 | 0.37653187 | 0.05954321 | 0.54977868 | 0.08592809 |
| rno-let-7a-5p | 28115 | 29099 | 18426 | 27691 | 0.41636407 | 0.04958619 | 0.60959713 | 0.07155242 |
| rno-let-7a-5p | 28105 | 29069 | 18434 | 27695 | 0.41560841 | 0.04841096 | 0.60845766 | 0.0698559 |
| rno-let-7c-5p | 27824 | 27785 | 18525 | 26665 | 0.40126001 | 0.04113866 | 0.5868563 | 0.05935891 |
| rno-let-7c-5p | 27661 | 27609 | 18433 | 26419 | 0.40039918 | 0.04405123 | 0.58556242 | 0.06356277 |
| rno-miR-191a-5p | 26357 | 24322 | 16659 | 20926 | 0.45090199 | 0.15010608 | 0.66188437 | 0.2169653 |
| rno-miR-29a-3p | 25661 | 27745 | 22461 | 23580 | 0.1329953 | 0.16229907 | 0.19215523 | 0.23466408 |
| rno-miR-378a-3p | 24555 | 21499 | 20531 | 23588 | 0.1785033 | -0.0926653 | 0.25821292 | -0.1337835 |
| rno-miR-23b-3p | 20242 | 20905 | 12637 | 15542 | 0.46260531 | 0.29429034 | 0.67969783 | 0.42767588 |
| rno-let-7d-5p | 19450 | 16710 | 11824 | 16185 | 0.48768945 | 0.03191974 | 0.71805198 | 0.04605437 |
| rno-miR-125a-5p | 19253 | 17243 | 14327 | 19076 | 0.29338892 | -0.1009389 | 0.42634672 | -0.1457479 |
| rno-miR-7a-5p | 18826 | 16987 | 12312 | 16555 | 0.41839553 | 0.02575875 | 0.61266136 | 0.03716408 |
| rno-miR-7a-5p | 18687 | 16809 | 12200 | 16411 | 0.42004727 | 0.02396147 | 0.61515383 | 0.03457075 |
| rno-miR-320-3p | 17715 | 17793 | 13065 | 16373 | 0.30214425 | 0.08312357 | 0.43926434 | 0.11999109 |
| rno-miR-93-5p | 15885 | 11953 | 9034 | 10695 | 0.54986155 | 0.11109149 | 0.81422827 | 0.16043627 |
| rno-miR-340-5p | 15756 | 15240 | 9455 | 13164 | 0.49986117 | 0.14617659 | 0.73675196 | 0.21126497 |
| rno-miR-340-5p | 15756 | 15240 | 9455 | 13164 | 0.49986117 | 0.14617659 | 0.73675196 | 0.21126497 |
| rno-miR-27a-3p | 15571 | 13800 | 10798 | 12947 | 0.362016 | 0.06378285 | 0.52809748 | 0.09205042 |
| rno-miR-25-3p | 14240 | 12138 | 10169 | 13157 | 0.33356549 | -0.0805693 | 0.48577133 | -0.1162998 |
| rno-miR-9a-5p | 14074 | 13247 | 9321 | 12691 | 0.40632614 | 0.04287146 | 0.59447577 | 0.06185992 |
| rno-miR-9a-5p | 14073 | 13247 | 9327 | 12693 | 0.40564103 | 0.04271396 | 0.59344488 | 0.06163258 |
| rno-miR-9a-5p | 14073 | 13247 | 9326 | 12693 | 0.40574384 | 0.04271396 | 0.59359957 | 0.06163258 |
| rno-miR-128-3p | 13685 | 12454 | 8718 | 11449 | 0.44342276 | 0.08408986 | 0.65052633 | 0.12138759 |
| rno-let-7b-5p | 13177 | 11801 | 8818 | 11614 | 0.39636281 | 0.01597267 | 0.57949857 | 0.02304418 |
| rno-miR-101a-3p | 12684 | 12845 | 9264 | 11300 | 0.31164571 | 0.12797681 | 0.45330262 | 0.18488412 |
| rno-miR-128-3p | 11661 | 10405 | 7174 | 9610 | 0.47645341 | 0.07944042 | 0.70084186 | 0.11466863 |
| rno-miR-26a-5p | 11420 | 15231 | 9186 | 10206 | 0.21683005 | 0.39509376 | 0.31405396 | 0.57759312 |
| rno-miR-30e-5p | 11408 | 10494 | 7150 | 9050 | 0.45888566 | 0.14776914 | 0.67403074 | 0.213575 |
| rno-miR-101b-3p | 11406 | 10803 | 7964 | 9904 | 0.35539494 | 0.08683054 | 0.51822781 | 0.12534879 |
| rno-miR-423-3p | 10487 | 9579 | 6707 | 9080 | 0.43968826 | 0.05348625 | 0.64486252 | 0.07718276 |
| rno-miR-328a-3p | 10307 | 8647 | 5978 | 7708 | 0.5316549 | 0.11482727 | 0.78588967 | 0.16584312 |
| rno-miR-152-3p | 9073 | 8472 | 6525 | 7967 | 0.32670855 | 0.06143926 | 0.47560176 | 0.08866601 |
| rno-miR-30c-5p | 8805 | 8162 | 6352 | 8054 | 0.32367883 | 0.01332018 | 0.47111209 | 0.01921724 |
| rno-miR-30c-5p | 8798 | 8157 | 6343 | 8047 | 0.32428505 | 0.01357689 | 0.47201026 | 0.01958762 |
| rno-miR-183-5p | 8227 | 9352 | 5289 | 7122 | 0.43474401 | 0.27072963 | 0.63737147 | 0.39299249 |
| rno-miR-185-5p | 8202 | 8538 | 5616 | 7244 | 0.3742944 | 0.16398429 | 0.54643281 | 0.23711162 |
| rno-miR-351-5p | 7985 | 7802 | 6029 | 7269 | 0.27914942 | 0.07073187 | 0.40537368 | 0.10208709 |
| rno-miR-351-5p | 7985 | 7802 | 6029 | 7269 | 0.27914942 | 0.07073187 | 0.40537368 | 0.10208709 |
| rno-let-7e-5p | 7883 | 8143 | 5306 | 7287 | 0.3907802 | 0.11095269 | 0.5711201 | 0.16023541 |
| rno-miR-344a-3p | 7789 | 7725 | 5210 | 7008 | 0.39679975 | 0.09733252 | 0.58015475 | 0.14053216 |
| rno-miR-344a-3p | 7789 | 7725 | 5210 | 7008 | 0.39679975 | 0.09733252 | 0.58015475 | 0.14053216 |
| rno-miR-210-3p | 7358 | 5324 | 4035 | 6322 | 0.58334065 | -0.1713893 | 0.866745 | -0.2478705 |
| rno-miR-140-3p | 7212 | 7467 | 5875 | 7019 | 0.2043249 | 0.06185282 | 0.29581055 | 0.08926323 |
| rno-miR-186-5p | 7181 | 6671 | 5151 | 5721 | 0.32922478 | 0.15332473 | 0.47933222 | 0.2216357 |
| rno-miR-182 | 6438 | 7009 | 4676 | 5627 | 0.31707756 | 0.21874011 | 0.46133765 | 0.31684266 |
| rno-miR-98-5p | 5494 | 5241 | 3447 | 4863 | 0.45789062 | 0.07482185 | 0.67251561 | 0.10799552 |
| rno-miR-92a-3p | 5412 | 4267 | 3119 | 4505 | 0.53756887 | -0.0542636 | 0.79507829 | -0.078305 |
| rno-miR-139-5p | 5169 | 5758 | 3089 | 4014 | 0.50375394 | 0.35693819 | 0.74274533 | 0.52052717 |
| rno-miR-20a-5p | 4908 | 4511 | 2809 | 3679 | 0.54399378 | 0.2031746 | 0.80507862 | 0.29413361 |
| rno-miR-148b-3p | 4454 | 3941 | 2800 | 3541 | 0.45602426 | 0.10692328 | 0.66967473 | 0.15440491 |
| rno-miR-16-5p | 4260 | 4149 | 3141 | 3826 | 0.30239157 | 0.08100313 | 0.43962949 | 0.11692678 |
| rno-miR-23a-3p | 4258 | 4889 | 3278 | 3428 | 0.26008493 | 0.3513286 | 0.3773601 | 0.5121723 |
| rno-miR-30b-5p | 4142 | 3818 | 2856 | 3129 | 0.36753358 | 0.198359 | 0.53633157 | 0.28711544 |
| rno-miR-151-3p | 3832 | 4035 | 2569 | 3128 | 0.39462584 | 0.25324585 | 0.57689067 | 0.36732816 |
| rno-miR-149-5p | 3819 | 4319 | 2201 | 2607 | 0.53754153 | 0.49436904 | 0.79503577 | 0.72830673 |
| rno-miR-374-5p | 3740 | 4040 | 2430 | 3120 | 0.42463533 | 0.25698324 | 0.62208196 | 0.37280926 |
| rno-miR-181d-5p | 3639 | 3308 | 2567 | 3261 | 0.34547212 | 0.01430964 | 0.50345876 | 0.02064479 |
| rno-miR-130b-5p | 3596 | 2912 | 1941 | 2954 | 0.59779664 | -0.0143198 | 0.8895929 | -0.0206595 |
| rno-miR-125b-5p | 3479 | 3939 | 2607 | 3433 | 0.28655932 | 0.13727618 | 0.4162821 | 0.19835956 |
| rno-miR-125b-5p | 3439 | 3849 | 2543 | 3345 | 0.29956536 | 0.14011676 | 0.43545765 | 0.20247746 |
| rno-miR-106b-3p | 3394 | 2808 | 2171 | 2837 | 0.43953279 | -0.0102746 | 0.64462684 | -0.0148232 |
| rno-miR-146b-5p | 3276 | 3808 | 2471 | 3078 | 0.28014616 | 0.2120244 | 0.40684035 | 0.30704025 |
| rno-miR-672-5p | 3258 | 2670 | 2350 | 3090 | 0.32382311 | -0.1458333 | 0.47132585 | -0.2107671 |
| rno-miR-322-3p | 3192 | 3400 | 2824 | 3178 | 0.12234043 | 0.06749772 | 0.17672056 | 0.09741562 |
| rno-miR-322-3p | 3192 | 3400 | 2824 | 3178 | 0.12234043 | 0.06749772 | 0.17672056 | 0.09741562 |
| rno-miR-598-3p | 3153 | 3411 | 1976 | 2705 | 0.45895886 | 0.23086985 | 0.67414222 | 0.33456616 |
| rno-miR-598-3p | 3153 | 3411 | 1976 | 2705 | 0.45895886 | 0.23086985 | 0.67414222 | 0.33456616 |
| rno-miR-298-5p | 3002 | 3326 | 1899 | 2404 | 0.45011222 | 0.32181501 | 0.66068407 | 0.46835127 |
| rno-miR-26b-5p | 2938 | 3182 | 2223 | 2519 | 0.27707809 | 0.23259077 | 0.40232645 | 0.33708271 |
| rno-miR-107-3p | 2879 | 2586 | 1812 | 2148 | 0.45491366 | 0.18504436 | 0.66798483 | 0.26772828 |
| rno-miR-378b | 2707 | 2326 | 2065 | 2439 | 0.26906957 | -0.0474292 | 0.39055311 | -0.0684387 |
| rno-miR-361-3p | 2577 | 2368 | 1522 | 2244 | 0.5147597 | 0.05377277 | 0.75972418 | 0.07759641 |
| rno-miR-450a-5p | 2533 | 2595 | 2004 | 2454 | 0.23319374 | 0.05585264 | 0.33796457 | 0.08059929 |
| rno-miR-872-5p | 2444 | 2390 | 1688 | 2145 | 0.36592449 | 0.10804851 | 0.53392938 | 0.15603297 |
| rno-miR-344b-3p | 2358 | 2050 | 1480 | 1921 | 0.45752996 | 0.06497104 | 0.67196654 | 0.09376639 |
| rno-miR-344b-3p | 2358 | 2050 | 1480 | 1921 | 0.45752996 | 0.06497104 | 0.67196654 | 0.09376639 |
| rno-miR-96-5p | 2267 | 2431 | 1707 | 1951 | 0.28183191 | 0.21907805 | 0.40932133 | 0.31733612 |
| rno-miR-106b-5p | 2219 | 1945 | 1422 | 1871 | 0.43779182 | 0.03878407 | 0.6419882 | 0.0559606 |
| rno-miR-423-5p | 2185 | 2009 | 1386 | 1845 | 0.4474937 | 0.08510638 | 0.65670602 | 0.12285675 |
| rno-miR-340-3p | 2072 | 1957 | 1444 | 1818 | 0.35722412 | 0.07364238 | 0.52095326 | 0.10629156 |
| rno-miR-340-3p | 2072 | 1957 | 1444 | 1818 | 0.35722412 | 0.07364238 | 0.52095326 | 0.10629156 |
| rno-miR-501-3p | 2050 | 1521 | 1165 | 1512 | 0.55054432 | 0.00593472 | 0.81529395 | 0.00856201 |
| rno-miR-484 | 1964 | 1840 | 1020 | 1522 | 0.63270777 | 0.18917311 | 0.94522578 | 0.27373741 |
| rno-miR-17-5p | 1922 | 1578 | 999 | 1264 | 0.63197535 | 0.22097115 | 0.94405175 | 0.32010074 |
| rno-let-7d-3p | 1856 | 1544 | 1146 | 1417 | 0.47301799 | 0.08578183 | 0.69558967 | 0.12383299 |
| rno-miR-1839-5p | 1843 | 1643 | 1147 | 1272 | 0.46555184 | 0.25454545 | 0.68419068 | 0.36923381 |
| rno-miR-17-5p | 1836 | 1516 | 959 | 1204 | 0.62754919 | 0.22941176 | 0.93696334 | 0.33243436 |
| rno-miR-29b-3p | 1728 | 1631 | 1201 | 1340 | 0.35984978 | 0.19589364 | 0.52486707 | 0.28352378 |
| rno-miR-29b-3p | 1728 | 1631 | 1201 | 1340 | 0.35984978 | 0.19589364 | 0.52486707 | 0.28352378 |
| rno-miR-29b-3p | 1726 | 1631 | 1202 | 1339 | 0.3579235 | 0.196633 | 0.52199557 | 0.28460082 |
| rno-miR-24-2-5p | 1712 | 2047 | 1332 | 1488 | 0.24967148 | 0.31626591 | 0.36208862 | 0.46013658 |
| rno-miR-99b-3p | 1658 | 1351 | 1005 | 1445 | 0.49042433 | -0.0672389 | 0.72224851 | -0.0970418 |
| rno-miR-425-5p | 1647 | 1515 | 965 | 1215 | 0.52220521 | 0.21978022 | 0.77123971 | 0.31836148 |
| rno-miR-450b-5p | 1537 | 1686 | 1298 | 1625 | 0.1686067 | 0.03684687 | 0.24382678 | 0.05316482 |
| rno-miR-450b-5p | 1537 | 1686 | 1298 | 1625 | 0.1686067 | 0.03684687 | 0.24382678 | 0.05316482 |
| rno-let-7a-1-3p | 1537 | 1238 | 1016 | 1346 | 0.40814728 | -0.0835913 | 0.59721676 | -0.1206671 |
| rno-let-7c-2-3p | 1536 | 1237 | 1016 | 1346 | 0.40752351 | -0.084398 | 0.59627781 | -0.1218329 |
| rno-miR-362-5p | 1527 | 988 | 719 | 1064 | 0.71950134 | -0.0740741 | 1.08663639 | -0.1069152 |
| rno-miR-195-5p | 1473 | 1510 | 1101 | 1269 | 0.28904429 | 0.17344368 | 0.41994296 | 0.25085648 |
| rno-miR-92a-3p | 1447 | 837 | 671 | 994 | 0.73276676 | -0.171491 | 1.10868025 | -0.2480182 |
| rno-miR-325-5p | 1443 | 1622 | 1092 | 1307 | 0.27692308 | 0.21509047 | 0.40209844 | 0.31151468 |
| rno-miR-192-5p | 1417 | 1452 | 1104 | 1410 | 0.24831416 | 0.0293501 | 0.36009959 | 0.04234629 |
| rno-miR-344b-1-3p | 1412 | 1474 | 811 | 1113 | 0.54071075 | 0.27908775 | 0.79996627 | 0.40528293 |
| rno-miR-143-3p | 1390 | 1347 | 1290 | 1570 | 0.07462687 | -0.1528968 | 0.10771382 | -0.2210147 |
| rno-miR-10a-5p | 1390 | 1459 | 1010 | 1492 | 0.31666667 | -0.0223653 | 0.46072959 | -0.0322677 |
| rno-miR-296-3p | 1388 | 1783 | 915 | 1222 | 0.41076856 | 0.3733777 | 0.60116392 | 0.54506242 |
| rno-miR-126a-3p | 1253 | 1253 | 965 | 1143 | 0.25969342 | 0.0918197 | 0.37678557 | 0.13256101 |
| rno-miR-140-5p | 1214 | 1123 | 904 | 1007 | 0.29272899 | 0.10892019 | 0.42537374 | 0.15729424 |
| rno-miR-384-5p | 1209 | 1286 | 838 | 1051 | 0.36248168 | 0.20111254 | 0.5287921 | 0.29112797 |
| rno-miR-195-3p | 1187 | 963 | 822 | 1156 | 0.36336486 | -0.1821614 | 0.53010964 | -0.2635337 |
| rno-miR-542-3p | 1130 | 1103 | 791 | 960 | 0.35294118 | 0.13863306 | 0.51457317 | 0.20032648 |
| rno-miR-542-3p | 1130 | 1103 | 791 | 960 | 0.35294118 | 0.13863306 | 0.51457317 | 0.20032648 |
| rno-miR-542-3p | 1130 | 1103 | 791 | 960 | 0.35294118 | 0.13863306 | 0.51457317 | 0.20032648 |
| rno-miR-503-5p | 1128 | 1176 | 1111 | 1137 | 0.01518535 | 0.03372244 | 0.02190825 | 0.04865581 |
| rno-miR-503-5p | 1128 | 1176 | 1111 | 1137 | 0.01518535 | 0.03372244 | 0.02190825 | 0.04865581 |
| rno-miR-384-3p | 1121 | 1178 | 729 | 899 | 0.42378378 | 0.26865672 | 0.62079556 | 0.38994652 |
| rno-miR-15b-5p | 1094 | 994 | 708 | 874 | 0.42841287 | 0.12847966 | 0.62779147 | 0.18561257 |
| rno-miR-361-5p | 1046 | 955 | 669 | 873 | 0.43965015 | 0.08971554 | 0.64480474 | 0.12951908 |
| rno-miR-181c-3p | 1008 | 937 | 712 | 870 | 0.34418605 | 0.07415606 | 0.50154649 | 0.10703365 |
| rno-miR-652-3p | 965 | 1230 | 557 | 712 | 0.53613666 | 0.53347065 | 0.79285161 | 0.78870917 |
| rno-miR-342-3p | 899 | 951 | 460 | 633 | 0.64606328 | 0.40151515 | 0.96668725 | 0.58723984 |
| rno-miR-325-3p | 892 | 846 | 576 | 686 | 0.43051771 | 0.20887728 | 0.6309749 | 0.30244909 |
| rno-miR-374-3p | 866 | 1095 | 576 | 946 | 0.40221914 | 0.14600686 | 0.58829821 | 0.21101878 |
| rno-miR-532-3p | 862 | 709 | 626 | 763 | 0.3172043 | -0.0733696 | 0.46152521 | -0.1058974 |
| rno-miR-322-5p | 825 | 790 | 713 | 757 | 0.14564369 | 0.04266322 | 0.21049204 | 0.06155935 |
| rno-miR-322-5p | 825 | 790 | 713 | 757 | 0.14564369 | 0.04266322 | 0.21049204 | 0.06155935 |
| rno-miR-203a-3p | 803 | 998 | 584 | 697 | 0.31578947 | 0.35516224 | 0.45943162 | 0.51788116 |
| rno-miR-335 | 801 | 851 | 545 | 637 | 0.38038633 | 0.28763441 | 0.55554601 | 0.41786576 |
| rno-miR-339-5p | 786 | 729 | 633 | 690 | 0.21564482 | 0.05496829 | 0.31232381 | 0.07932245 |
| rno-miR-421-3p | 785 | 836 | 494 | 563 | 0.455043 | 0.39027877 | 0.66818161 | 0.57036802 |
| rno-miR-499-5p | 745 | 872 | 610 | 738 | 0.19926199 | 0.16645963 | 0.28843118 | 0.24070732 |
| rno-miR-3559-5p | 724 | 761 | 493 | 685 | 0.37962202 | 0.10511757 | 0.55440205 | 0.15179247 |
| rno-miR-132-3p | 684 | 1135 | 562 | 675 | 0.19582665 | 0.50828729 | 0.28342619 | 0.74973289 |
| rno-miR-674-3p | 645 | 695 | 430 | 572 | 0.4 | 0.19415943 | 0.5849625 | 0.28099783 |
| rno-miR-301a-5p | 636 | 487 | 386 | 496 | 0.48923679 | -0.0183113 | 0.72042592 | -0.0264183 |
| rno-miR-378a-5p | 614 | 520 | 458 | 547 | 0.29104478 | -0.0506092 | 0.42289106 | -0.0730292 |
| rno-miR-702-3p | 612 | 534 | 294 | 430 | 0.70198675 | 0.21576763 | 1.0577155 | 0.31250308 |
| rno-miR-29c-3p | 608 | 601 | 453 | 496 | 0.29217719 | 0.19143118 | 0.42456027 | 0.27702487 |
| rno-miR-29c-3p | 608 | 601 | 453 | 496 | 0.29217719 | 0.19143118 | 0.42456027 | 0.27702487 |
| rno-miR-760-3p | 587 | 965 | 335 | 472 | 0.54663774 | 0.6861517 | 0.80919941 | 1.03174208 |
| rno-miR-362-3p | 574 | 462 | 476 | 524 | 0.18666667 | -0.1257606 | 0.27008916 | -0.181674 |
| rno-miR-181c-5p | 560 | 467 | 474 | 512 | 0.16634429 | -0.0919305 | 0.24053977 | -0.1327213 |
| rno-miR-19b-3p | 546 | 439 | 393 | 494 | 0.32587859 | -0.1178992 | 0.47437164 | -0.1702901 |
| rno-miR-19b-3p | 546 | 442 | 393 | 495 | 0.32587859 | -0.113127 | 0.47437164 | -0.1633822 |
| rno-miR-330-5p | 546 | 509 | 367 | 460 | 0.39211391 | 0.10113519 | 0.57312089 | 0.1460318 |
| rno-miR-296-5p | 531 | 463 | 285 | 394 | 0.60294118 | 0.16102684 | 0.89774994 | 0.23281656 |
| rno-miR-27b-5p | 528 | 624 | 502 | 560 | 0.05048544 | 0.10810811 | 0.07285057 | 0.1561192 |
| rno-miR-15a-5p | 523 | 454 | 328 | 426 | 0.45828437 | 0.06363636 | 0.67311513 | 0.09183887 |
| rno-miR-132-5p | 492 | 713 | 444 | 504 | 0.1025641 | 0.34346754 | 0.14809864 | 0.50047834 |
| rno-miR-1843a-5p | 488 | 483 | 315 | 447 | 0.43088418 | 0.07741935 | 0.63152932 | 0.11174836 |
| rno-miR-30a-3p | 484 | 439 | 361 | 483 | 0.29112426 | -0.0954447 | 0.42300821 | -0.1378022 |
| rno-miR-15b-3p | 471 | 358 | 253 | 312 | 0.60220994 | 0.13731343 | 0.89658967 | 0.19841356 |
| rno-miR-130a-3p | 467 | 433 | 336 | 368 | 0.32627646 | 0.16229713 | 0.47496132 | 0.23466126 |
| rno-miR-30c-2-3p | 453 | 388 | 290 | 398 | 0.43876178 | -0.0254453 | 0.64345815 | -0.0367118 |
| rno-miR-24-1-5p | 440 | 424 | 318 | 368 | 0.32189974 | 0.14141414 | 0.46847676 | 0.2043585 |
| rno-miR-505-3p | 438 | 346 | 195 | 280 | 0.76777251 | 0.21086262 | 1.16745675 | 0.30534521 |
| rno-miR-152-5p | 432 | 416 | 281 | 449 | 0.42356241 | -0.0763006 | 0.62046118 | -0.1101319 |
| rno-miR-365-3p | 414 | 452 | 207 | 275 | 0.66666667 | 0.4869326 | 1 | 0.71689115 |
| rno-let-7i-3p | 397 | 267 | 239 | 286 | 0.49685535 | -0.0687161 | 0.73212839 | -0.0991754 |
| rno-miR-10b-3p | 371 | 376 | 242 | 317 | 0.42088091 | 0.17027417 | 0.61641214 | 0.24624982 |
| rno-miR-324-3p | 355 | 283 | 209 | 273 | 0.5177305 | 0.03597122 | 0.76431608 | 0.0519011 |
| rno-miR-32-5p | 354 | 370 | 244 | 309 | 0.36789298 | 0.17967599 | 0.53686821 | 0.25991843 |
| rno-miR-497-5p | 353 | 337 | 262 | 266 | 0.29593496 | 0.23548922 | 0.43010137 | 0.34132235 |
| rno-miR-872-3p | 351 | 288 | 215 | 269 | 0.48056537 | 0.06822262 | 0.70713437 | 0.09846264 |
| rno-miR-330-3p | 350 | 300 | 248 | 287 | 0.34113712 | 0.04429302 | 0.4970148 | 0.06391176 |
| rno-miR-193b-3p | 339 | 428 | 290 | 295 | 0.15580286 | 0.36791148 | 0.22523237 | 0.53689584 |
| rno-miR-151-5p | 324 | 290 | 207 | 241 | 0.44067797 | 0.18455744 | 0.64636305 | 0.26701975 |
| rno-miR-147 | 315 | 330 | 236 | 278 | 0.28675136 | 0.17105263 | 0.41656497 | 0.24738114 |
| rno-miR-30d-3p | 311 | 284 | 237 | 251 | 0.27007299 | 0.12336449 | 0.39202752 | 0.17820357 |
| rno-miR-27a-5p | 307 | 380 | 372 | 370 | 0.19145803 | 0.02666667 | -0.277064 | 0.03847415 |
| rno-miR-129-5p | 306 | 276 | 214 | 307 | 0.35384615 | -0.1063465 | 0.51592086 | -0.1535704 |
| rno-miR-129-5p | 303 | 280 | 212 | 301 | 0.35339806 | -0.0722892 | 0.51525353 | -0.1043367 |
| rno-miR-181b-5p | 302 | 242 | 154 | 204 | 0.64912281 | 0.17040359 | 0.9716182 | 0.2464379 |
| rno-miR-29c-5p | 293 | 292 | 229 | 243 | 0.24521073 | 0.18317757 | 0.35555307 | 0.26501206 |
| rno-miR-29c-5p | 293 | 292 | 229 | 243 | 0.24521073 | 0.18317757 | 0.35555307 | 0.26501206 |
| rno-miR-3068-3p | 293 | 267 | 175 | 230 | 0.5042735 | 0.14889336 | 0.74354574 | 0.21520588 |
| rno-miR-139-3p | 292 | 289 | 162 | 225 | 0.57268722 | 0.24902724 | 0.84997456 | 0.36114449 |
| rno-miR-23b-5p | 290 | 249 | 219 | 312 | 0.27897839 | -0.2245989 | 0.40512203 | -0.3254003 |
| rno-miR-188-5p | 279 | 161 | 123 | 206 | 0.7761194 | -0.2452316 | 1.18160681 | -0.3555836 |
| rno-miR-22-5p | 273 | 325 | 264 | 244 | 0.03351955 | 0.28471002 | 0.04836302 | 0.41355857 |
| rno-miR-212-5p | 263 | 420 | 243 | 307 | 0.07905138 | 0.31086657 | 0.11410649 | 0.45215067 |
| rno-miR-674-5p | 261 | 260 | 169 | 213 | 0.42790698 | 0.1987315 | 0.62702656 | 0.28765819 |
| rno-miR-1224 | 257 | 298 | 234 | 216 | 0.09368635 | 0.31906615 | 0.13525983 | 0.46428102 |
| rno-miR-488-3p | 251 | 257 | 200 | 216 | 0.22616408 | 0.17336152 | 0.32768736 | 0.25073705 |
| rno-miR-331-3p | 251 | 244 | 155 | 213 | 0.4729064 | 0.1356674 | 0.69541915 | 0.19602772 |
| rno-miR-344b-5p | 250 | 245 | 157 | 221 | 0.45700246 | 0.10300429 | 0.67116354 | 0.14873538 |
| rno-miR-344b-5p | 250 | 245 | 157 | 221 | 0.45700246 | 0.10300429 | 0.67116354 | 0.14873538 |
| rno-miR-344b-5p | 250 | 245 | 157 | 221 | 0.45700246 | 0.10300429 | 0.67116354 | 0.14873538 |
| rno-miR-30e-3p | 248 | 305 | 201 | 266 | 0.20935412 | 0.13660245 | 0.30314462 | 0.197383 |
| rno-miR-190a-5p | 246 | 195 | 162 | 145 | 0.41176471 | 0.29411765 | 0.6026645 | 0.42742122 |
| rno-miR-190a-5p | 246 | 195 | 162 | 145 | 0.41176471 | 0.29411765 | 0.6026645 | 0.42742122 |
| rno-miR-671 | 245 | 286 | 177 | 253 | 0.32227488 | 0.12244898 | 0.46903239 | 0.17687776 |
| rno-miR-879-5p | 238 | 201 | 152 | 179 | 0.44102564 | 0.11578947 | 0.64689025 | 0.16723591 |
| rno-miR-1843b-5p | 223 | 197 | 133 | 197 | 0.50561798 | 0 | 0.74561746 | 0 |
| rno-miR-29b-5p | 221 | 196 | 129 | 164 | 0.52571429 | 0.17777778 | 0.7766753 | 0.25715784 |
| rno-miR-29b-5p | 221 | 196 | 129 | 164 | 0.52571429 | 0.17777778 | 0.7766753 | 0.25715784 |
| rno-miR-326-3p | 213 | 261 | 121 | 136 | 0.5508982 | 0.62972292 | 0.81584638 | 0.94044316 |
| rno-miR-3559-3p | 212 | 171 | 172 | 185 | 0.20833333 | -0.0786517 | 0.3016557 | -0.1135289 |
| rno-miR-30b-3p | 208 | 205 | 217 | 250 | 0.04235294 | -0.1978022 | -0.0611115 | -0.2863042 |
| rno-miR-301a-3p | 204 | 186 | 110 | 151 | 0.59872611 | 0.20771513 | 0.89106563 | 0.30075407 |
| rno-miR-130b-3p | 203 | 161 | 108 | 130 | 0.61093248 | 0.21305842 | 0.91044842 | 0.30854907 |
| rno-miR-7b | 202 | 225 | 123 | 142 | 0.48615385 | 0.45231608 | 0.71569698 | 0.66403407 |
| rno-miR-93-3p | 202 | 151 | 118 | 161 | 0.525 | -0.0641026 | 0.77556843 | -0.0925121 |
| rno-miR-148b-5p | 198 | 199 | 140 | 179 | 0.34319527 | 0.10582011 | 0.5000736 | 0.15280884 |
| rno-miR-9a-3p | 188 | 200 | 143 | 171 | 0.27190332 | 0.15633423 | 0.39471751 | 0.22600367 |
| rno-miR-9a-3p | 188 | 200 | 143 | 171 | 0.27190332 | 0.15633423 | 0.39471751 | 0.22600367 |
| rno-miR-9a-3p | 185 | 191 | 141 | 168 | 0.26993865 | 0.1281337 | 0.39183011 | 0.18511141 |
| rno-miR-324-5p | 184 | 190 | 139 | 124 | 0.27863777 | 0.42038217 | 0.40462088 | 0.6156593 |
| rno-miR-1249 | 183 | 206 | 92 | 110 | 0.66181818 | 0.60759494 | 0.99213788 | 0.90514081 |
| rno-miR-298-3p | 181 | 197 | 111 | 166 | 0.47945205 | 0.1707989 | 0.70543002 | 0.24701239 |
| rno-miR-500-3p | 175 | 151 | 141 | 140 | 0.21518987 | 0.07560137 | 0.31165976 | 0.10912172 |
| rno-miR-350 | 173 | 213 | 96 | 122 | 0.57249071 | 0.54328358 | 0.84966573 | 0.80397228 |
| rno-miR-350 | 173 | 213 | 96 | 122 | 0.57249071 | 0.54328358 | 0.84966573 | 0.80397228 |
| rno-miR-877 | 171 | 111 | 78 | 103 | 0.74698795 | 0.07476636 | 1.1324503 | 0.10791534 |
| rno-miR-125b-1-3p | 167 | 202 | 135 | 189 | 0.21192053 | 0.06649616 | 0.3068887 | 0.09596906 |
| rno-miR-194-5p | 160 | 155 | 154 | 170 | 0.03821656 | -0.0923077 | 0.05514155 | -0.1332665 |
| rno-miR-3068-5p | 160 | 152 | 110 | 150 | 0.37037037 | 0.01324503 | 0.54056838 | 0.01910882 |
| rno-miR-25-5p | 160 | 103 | 97 | 117 | 0.49027237 | -0.1272727 | 0.72201525 | -0.1838642 |
| rno-miR-138-5p | 157 | 138 | 93 | 70 | 0.512 | 0.65384615 | 0.75546194 | 0.97924144 |
| rno-miR-339-3p | 153 | 164 | 150 | 128 | 0.01980198 | 0.24657534 | 0.02856915 | 0.357552 |
| rno-miR-148a-3p | 149 | 150 | 104 | 147 | 0.35573123 | 0.02020202 | 0.5187288 | 0.02914635 |
| rno-miR-18a-5p | 148 | 144 | 76 | 79 | 0.64285714 | 0.58295964 | 0.96152585 | 0.86614425 |
| rno-miR-33-5p | 146 | 213 | 94 | 132 | 0.43333333 | 0.46956522 | 0.63523571 | 0.6903155 |
| rno-miR-3473 | 143 | 160 | 119 | 150 | 0.18320611 | 0.06451613 | 0.26505357 | 0.0931094 |
| rno-let-7f-2-3p | 142 | 131 | 96 | 130 | 0.38655462 | 0.00766284 | 0.56478462 | 0.01105519 |
| rno-miR-301b-5p | 142 | 106 | 64 | 117 | 0.75728155 | -0.0986547 | 1.14974712 | -0.1424443 |
| rno-miR-338-5p | 138 | 131 | 106 | 129 | 0.26229508 | 0.01538462 | 0.380604 | 0.02219575 |
| rno-let-7b-3p | 132 | 133 | 95 | 127 | 0.32599119 | 0.04615385 | 0.47453851 | 0.06659775 |
| rno-miR-194-5p | 126 | 107 | 105 | 116 | 0.18181818 | -0.0807175 | 0.26303441 | -0.116514 |
| rno-miR-138-5p | 126 | 102 | 77 | 54 | 0.48275862 | 0.61538462 | 0.71049338 | 0.91753784 |
| rno-miR-1306-5p | 125 | 124 | 84 | 93 | 0.3923445 | 0.28571429 | 0.57346686 | 0.4150375 |
| rno-miR-503-3p | 114 | 106 | 116 | 142 | 0.0173913 | -0.2903226 | -0.025091 | -0.4218267 |
| rno-miR-503-3p | 114 | 106 | 116 | 142 | 0.0173913 | -0.2903226 | -0.025091 | -0.4218267 |
| rno-miR-6329 | 106 | 90 | 87 | 90 | 0.19689119 | 0 | 0.28497696 | 0 |
| rno-let-7e-3p | 104 | 111 | 71 | 115 | 0.37714286 | -0.0353982 | 0.5506926 | -0.0510742 |
| rno-miR-19a-3p | 104 | 86 | 57 | 80 | 0.58385093 | 0.07228916 | 0.8675497 | 0.10433666 |
| rno-miR-450b-3p | 98 | 117 | 96 | 127 | 0.02061856 | -0.0819672 | 0.02974734 | -0.11832 |
| rno-miR-450b-3p | 98 | 117 | 96 | 127 | 0.02061856 | -0.0819672 | 0.02974734 | -0.11832 |
| rno-miR-150-5p | 96 | 104 | 92 | 87 | 0.04255319 | 0.17801047 | 0.06140054 | 0.25749622 |
| rno-miR-1843a-3p | 96 | 105 | 66 | 75 | 0.37037037 | 0.33333333 | 0.54056838 | 0.48542683 |
| rno-miR-218a-2-3p | 89 | 82 | 57 | 57 | 0.43835616 | 0.35971223 | 0.64284342 | 0.52466199 |
| rno-miR-425-3p | 84 | 77 | 51 | 66 | 0.48888889 | 0.15384615 | 0.71989208 | 0.22239242 |
| rno-miR-455-5p | 83 | 121 | 68 | 97 | 0.1986755 | 0.22018349 | 0.28757659 | 0.3189504 |
| rno-miR-126a-5p | 83 | 80 | 64 | 61 | 0.2585034 | 0.26950355 | 0.37503943 | 0.39119076 |
| rno-miR-342-5p | 82 | 75 | 58 | 82 | 0.34285714 | -0.089172 | 0.49957101 | -0.1287333 |
| rno-miR-1306-3p | 79 | 61 | 28 | 54 | 0.95327103 | 0.12173913 | 1.49642583 | 0.17584984 |
| rno-miR-3099 | 78 | 74 | 41 | 65 | 0.62184874 | 0.1294964 | 0.92785021 | 0.18708555 |
| rno-miR-125a-3p | 77 | 106 | 93 | 113 | 0.18823529 | -0.0639269 | -0.2723723 | -0.0922585 |
| rno-miR-345-3p | 75 | 65 | 50 | 57 | 0.4 | 0.13114754 | 0.5849625 | 0.1894778 |
| rno-miR-331-5p | 74 | 86 | 79 | 73 | 0.06535948 | 0.16352201 | -0.0943274 | 0.2364402 |
| rno-miR-191a-3p | 73 | 77 | 39 | 72 | 0.60714286 | 0.06711409 | 0.90442234 | 0.09686154 |
| rno-miR-615 | 71 | 66 | 51 | 68 | 0.32786885 | -0.0298507 | 0.47732178 | -0.0430687 |
| rno-miR-26b-3p | 71 | 56 | 39 | 50 | 0.58181818 | 0.11320755 | 0.8643449 | 0.16349873 |
| rno-miR-137-3p | 69 | 99 | 54 | 53 | 0.24390244 | 0.60526316 | 0.35363695 | 0.90143617 |
| rno-miR-92a-1-5p | 68 | 56 | 86 | 74 | 0.23376623 | -0.2769231 | -0.3388019 | -0.4020984 |
| rno-let-7f-1-3p | 68 | 51 | 44 | 62 | 0.42857143 | -0.1946903 | 0.62803122 | -0.281771 |
| rno-miR-210-5p | 67 | 39 | 38 | 77 | 0.55238095 | -0.6551724 | 0.81816168 | -0.9813843 |
| rno-miR-219a-1-3p | 65 | 64 | 52 | 70 | 0.22222222 | -0.0895522 | 0.32192809 | -0.129283 |
| rno-miR-672-3p | 62 | 58 | 46 | 58 | 0.2962963 | 0 | 0.43063435 | 0 |
| rno-miR-664-3p | 51 | 54 | 30 | 38 | 0.51851852 | 0.34782609 | 0.76553475 | 0.50695999 |
| rno-miR-664-3p | 51 | 54 | 30 | 38 | 0.51851852 | 0.34782609 | 0.76553475 | 0.50695999 |
| rno-miR-293-5p | 50 | 33 | 36 | 36 | 0.3255814 | -0.0869565 | 0.47393119 | -0.1255309 |
| rno-miR-212-3p | 49 | 110 | 52 | 60 | 0.05940594 | 0.58823529 | -0.0857299 | 0.87446912 |
| rno-miR-7a-1-3p | 49 | 71 | 41 | 44 | 0.17777778 | 0.46956522 | 0.25715784 | 0.6903155 |
| rno-miR-17-1-3p | 48 | 51 | 31 | 26 | 0.43037975 | 0.64935065 | 0.63076619 | 0.97198562 |
| rno-miR-200c-3p | 48 | 35 | 20 | 21 | 0.82352941 | 0.5 | 1.26303441 | 0.73696559 |
| rno-miR-34c-5p | 46 | 25 | 48 | 38 | 0.04255319 | -0.4126984 | -0.0614005 | -0.6040713 |
| rno-miR-219a-5p | 46 | 24 | 29 | 31 | 0.45333333 | -0.2545455 | 0.66558096 | -0.3692338 |
| rno-miR-153-3p | 45 | 44 | 39 | 37 | 0.14285714 | 0.17283951 | 0.20645088 | 0.24997825 |
| rno-miR-128-1-5p | 44 | 37 | 35 | 23 | 0.2278481 | 0.46666667 | 0.3301486 | 0.68589141 |
| rno-miR-183-3p | 43 | 43 | 26 | 37 | 0.49275362 | 0.15 | 0.72582504 | 0.21681139 |
| rno-miR-676 | 42 | 42 | 27 | 28 | 0.43478261 | 0.4 | 0.63742992 | 0.5849625 |
| rno-miR-184 | 40 | 81 | 23 | 25 | 0.53968254 | 1.05660377 | 0.79836614 | 1.69599381 |
| rno-miR-1949 | 39 | 55 | 53 | 63 | 0.30434783 | -0.1355932 | -0.4425182 | -0.1959202 |
| rno-miR-193a-5p | 37 | 31 | 30 | 39 | 0.20895522 | -0.2285714 | 0.30256277 | -0.3312059 |
| rno-miR-127-3p | 37 | 84 | 24 | 51 | 0.42622951 | 0.48888889 | 0.62449086 | 0.71989208 |
| rno-miR-874-3p | 36 | 56 | 35 | 39 | 0.02816901 | 0.35789474 | 0.04064198 | 0.5219527 |
| rno-miR-16-3p | 36 | 30 | 18 | 30 | 0.66666667 | 0 | 1 | 0 |
| rno-miR-196a-5p | 35 | 42 | 27 | 33 | 0.25806452 | 0.24 | 0.37439551 | 0.3479233 |
| rno-miR-1839-3p | 35 | 32 | 22 | 19 | 0.45614035 | 0.50980392 | 0.6698514 | 0.75207249 |
| rno-miR-100-5p | 34 | 48 | 26 | 38 | 0.26666667 | 0.23255814 | 0.38702312 | 0.33703499 |
| rno-miR-29a-5p | 33 | 25 | 23 | 16 | 0.35714286 | 0.43902439 | 0.52083216 | 0.64385619 |
| rno-miR-411-5p | 33 | 59 | 23 | 28 | 0.35714286 | 0.71264368 | 0.52083216 | 1.07528813 |
| rno-miR-341 | 33 | 37 | 5 | 28 | 1.47368421 | 0.27692308 | 2.72246602 | 0.40209844 |
| rno-miR-129-1-3p | 32 | 55 | 21 | 27 | 0.41509434 | 0.68292683 | 0.60768258 | 1.02647221 |
| rno-miR-379-5p | 32 | 66 | 20 | 26 | 0.46153846 | 0.86956522 | 0.67807191 | 1.3439544 |
| rno-miR-338-3p | 31 | 23 | 24 | 33 | 0.25454545 | -0.3571429 | 0.36923381 | -0.5208322 |
| rno-miR-30c-1-3p | 30 | 24 | 15 | 28 | 0.66666667 | -0.1538462 | 1 | -0.2223924 |
| rno-miR-98-3p | 29 | 28 | 18 | 26 | 0.46808511 | 0.07407407 | 0.68805599 | 0.1069152 |
| rno-miR-505-5p | 29 | 20 | 15 | 20 | 0.63636364 | 0 | 0.9510904 | 0 |
| rno-miR-181b-5p | 29 | 20 | 10 | 14 | 0.97435897 | 0.35294118 | 1.5360529 | 0.51457317 |
| rno-miR-18a-3p | 28 | 26 | 21 | 22 | 0.28571429 | 0.16666667 | 0.4150375 | 0.2410081 |
| rno-miR-6319 | 28 | 24 | 15 | 24 | 0.60465116 | 0 | 0.90046433 | 0 |
| rno-miR-6319 | 28 | 24 | 15 | 24 | 0.60465116 | 0 | 0.90046433 | 0 |
| rno-miR-450a-3p | 28 | 23 | 13 | 21 | 0.73170732 | 0.09090909 | 1.1069152 | 0.13124453 |
| rno-miR-6318 | 26 | 5 | 9 | 6 | 0.97142857 | -0.1818182 | 1.53051472 | -0.2630344 |
| rno-miR-92b-3p | 26 | 11 | 5 | 16 | 1.35483871 | -0.3703704 | 2.37851162 | -0.5405684 |
| rno-miR-345-5p | 25 | 12 | 22 | 16 | 0.12765957 | -0.2857143 | 0.18442457 | -0.4150375 |
| rno-miR-541-5p | 25 | 110 | 20 | 35 | 0.22222222 | 1.03448276 | 0.32192809 | 1.6520767 |
| rno-miR-501-5p | 25 | 28 | 17 | 36 | 0.38095238 | -0.25 | 0.55639335 | -0.3625701 |
| rno-miR-351-3p | 24 | 30 | 20 | 26 | 0.18181818 | 0.14285714 | 0.26303441 | 0.20645088 |
| rno-miR-351-3p | 24 | 30 | 20 | 26 | 0.18181818 | 0.14285714 | 0.26303441 | 0.20645088 |
| rno-miR-3064-5p | 24 | 16 | 7 | 17 | 1.09677419 | -0.0606061 | 1.77760758 | -0.0874628 |
| rno-miR-193b-5p | 23 | 21 | 21 | 10 | 0.09090909 | 0.70967742 | 0.13124453 | 1.07038933 |
| rno-miR-3102 | 23 | 24 | 16 | 18 | 0.35897436 | 0.28571429 | 0.52356196 | 0.4150375 |
| rno-miR-664-2-5p | 23 | 20 | 13 | 20 | 0.55555556 | 0 | 0.82312224 | 0 |
| rno-miR-34a-5p | 20 | 13 | 25 | 20 | 0.22222222 | -0.4242424 | -0.3219281 | -0.6214884 |
| rno-miR-145-5p | 19 | 22 | 17 | 24 | 0.11111111 | -0.0869565 | 0.16046467 | -0.1255309 |
| rno-miR-6315 | 18 | 12 | 8 | 13 | 0.76923077 | -0.08 | 1.169925 | -0.1154772 |
| rno-miR-130a-5p | 17 | 31 | 27 | 17 | 0.45454545 | 0.58333333 | -0.6674247 | 0.86673347 |
| rno-miR-199a-3p | 17 | 25 | 23 | 21 | 0.3 | 0.17391304 | -0.4360991 | 0.25153877 |
| rno-miR-200a-3p | 17 | 10 | 20 | 19 | 0.16216216 | -0.6206897 | -0.2344653 | -0.9259994 |
| rno-miR-301b-3p | 16 | 20 | 15 | 18 | 0.06451613 | 0.10526316 | 0.0931094 | 0.15200309 |
| rno-miR-129-1-3p | 16 | 19 | 12 | 14 | 0.28571429 | 0.3030303 | 0.4150375 | 0.44057259 |
| rno-miR-129-2-3p | 16 | 19 | 12 | 14 | 0.28571429 | 0.3030303 | 0.4150375 | 0.44057259 |
| rno-miR-146b-3p | 16 | 6 | 10 | 9 | 0.46153846 | -0.4 | 0.67807191 | -0.5849625 |
| rno-miR-879-3p | 16 | 23 | 9 | 26 | 0.56 | -0.122449 | 0.830075 | -0.1768778 |
| rno-miR-3577 | 16 | 12 | 5 | 12 | 1.04761905 | 0 | 1.67807191 | 0 |
| rno-miR-218a-1-3p | 15 | 17 | 24 | 26 | 0.46153846 | -0.4186047 | -0.6780719 | -0.6129769 |
| rno-miR-190b-5p | 15 | 14 | 10 | 13 | 0.4 | 0.07407407 | 0.5849625 | 0.1069152 |
| rno-miR-421-5p | 15 | 13 | 7 | 16 | 0.72727273 | -0.2068966 | 1.09953567 | -0.2995603 |
| rno-miR-3074 | 15 | 9 | 6 | 5 | 0.85714286 | 0.57142857 | 1.32192809 | 0.84799691 |
| rno-let-7g-3p | 15 | 14 | 3 | 6 | 1.33333333 | 0.8 | 2.32192809 | 1.22239242 |
| rno-miR-702-5p | 14 | 13 | 19 | 11 | 0.3030303 | 0.16666667 | -0.4405726 | 0.2410081 |
| rno-miR-486 | 14 | 10 | 11 | 8 | 0.24 | 0.22222222 | 0.3479233 | 0.32192809 |
| rno-miR-20b-5p | 14 | 10 | 5 | 5 | 0.94736842 | 0.66666667 | 1.48542683 | 1 |
| rno-miR-500-5p | 14 | 14 | 4 | 7 | 1.11111111 | 0.66666667 | 1.80735492 | 1 |
| rno-miR-382-3p | 14 | 42 | 2 | 6 | 1.5 | 1.5 | 2.80735492 | 2.80735492 |
| rno-miR-344i | 13 | 12 | 8 | 14 | 0.47619048 | -0.1538462 | 0.70043972 | -0.2223924 |
| rno-miR-145-3p | 12 | 9 | 8 | 10 | 0.4 | -0.1052632 | 0.5849625 | -0.1520031 |
| rno-miR-3075 | 12 | 14 | 7 | 8 | 0.52631579 | 0.54545455 | 0.77760758 | 0.80735492 |
| rno-miR-370-3p | 12 | 16 | 6 | 18 | 0.66666667 | -0.1176471 | 1 | -0.169925 |
| rno-miR-148a-5p | 12 | 13 | 6 | 12 | 0.66666667 | 0.08 | 1 | 0.11547722 |
| rno-miR-434-5p | 12 | 23 | 4 | 8 | 1 | 0.96774194 | 1.5849625 | 1.52356196 |
| rno-miR-434-5p | 12 | 23 | 4 | 8 | 1 | 0.96774194 | 1.5849625 | 1.52356196 |
| rno-miR-1956-5p | 11 | 24 | 18 | 17 | 0.48275862 | 0.34146341 | -0.7104934 | 0.49749966 |
| rno-miR-3084a-3p | 11 | 7 | 7 | 4 | 0.44444444 | 0.54545455 | 0.6520767 | 0.80735492 |
| rno-miR-3084b-3p | 11 | 7 | 7 | 4 | 0.44444444 | 0.54545455 | 0.6520767 | 0.80735492 |
| rno-miR-3084b-3p | 11 | 7 | 7 | 4 | 0.44444444 | 0.54545455 | 0.6520767 | 0.80735492 |
| rno-miR-3084d | 11 | 7 | 7 | 4 | 0.44444444 | 0.54545455 | 0.6520767 | 0.80735492 |
| rno-miR-433-3p | 11 | 19 | 7 | 9 | 0.44444444 | 0.71428571 | 0.6520767 | 1.07800251 |
| rno-miR-409a-3p | 11 | 34 | 5 | 10 | 0.75 | 1.09090909 | 1.13750352 | 1.76553475 |
| rno-miR-32-3p | 11 | 4 | 4 | 10 | 0.93333333 | -0.8571429 | 1.45943162 | -1.3219281 |
| rno-miR-455-3p | 11 | 10 | 3 | 7 | 1.14285714 | 0.35294118 | 1.87446912 | 0.51457317 |
| rno-miR-542-5p | 10 | 23 | 15 | 23 | 0.4 | 0 | -0.5849625 | 0 |
| rno-miR-542-5p | 10 | 23 | 15 | 23 | 0.4 | 0 | -0.5849625 | 0 |
| rno-miR-542-5p | 10 | 23 | 15 | 23 | 0.4 | 0 | -0.5849625 | 0 |
| rno-miR-434-3p | 10 | 12 | 6 | 9 | 0.5 | 0.28571429 | 0.73696559 | 0.4150375 |
| rno-miR-434-3p | 10 | 12 | 6 | 9 | 0.5 | 0.28571429 | 0.73696559 | 0.4150375 |
| rno-miR-99a-5p | 9 | 7 | 12 | 10 | 0.28571429 | -0.3529412 | -0.4150375 | -0.5145732 |
| rno-miR-200b-3p | 9 | 4 | 9 | 13 | 0 | -1.0588235 | 0 | -1.7004397 |
| rno-miR-221-3p | 9 | 63 | 7 | 12 | 0.25 | 1.36 | 0.36257008 | 2.39231742 |
| rno-miR-1956-3p | 9 | 10 | 4 | 11 | 0.76923077 | -0.0952381 | 1.169925 | -0.1375035 |
| rno-miR-196c-5p | 9 | 11 | 2 | 2 | 1.27272727 | 1.38461538 | 2.169925 | 2.45943162 |
| rno-miR-741-3p | 8 | 5 | 11 | 10 | 0.31578947 | -0.6666667 | -0.4594316 | -1 |
| rno-miR-190a-3p | 8 | 6 | 11 | 6 | 0.31578947 | 0 | -0.4594316 | 0 |
| rno-miR-190a-3p | 8 | 6 | 11 | 6 | 0.31578947 | 0 | -0.4594316 | 0 |
| rno-miR-494-3p | 7 | 14 | 10 | 5 | 0.35294118 | 0.94736842 | -0.5145732 | 1.48542683 |
| rno-miR-328b-3p | 7 | 14 | 7 | 14 | 0 | 0 | 0 | 0 |
| rno-miR-300-3p | 7 | 37 | 5 | 9 | 0.33333333 | 1.2173913 | 0.48542683 | 2.03952836 |
| rno-miR-323-3p | 7 | 18 | 3 | 8 | 0.8 | 0.76923077 | 1.22239242 | 1.169925 |
| rno-miR-5132-5p | 6 | 9 | 15 | 7 | 0.85714286 | 0.25 | -1.3219281 | 0.36257008 |
| rno-miR-382-5p | 6 | 34 | 9 | 9 | 0.4 | 1.1627907 | -0.5849625 | 1.91753784 |
| rno-miR-487b-3p | 6 | 19 | 5 | 4 | 0.18181818 | 1.30434783 | 0.26303441 | 2.24792751 |
| rno-miR-134-5p | 6 | 29 | 3 | 8 | 0.66666667 | 1.13513514 | 1 | 1.857981 |
| rno-miR-409a-5p | 6 | 17 | 2 | 10 | 1 | 0.51851852 | 1.5849625 | 0.76553475 |
| rno-miR-543-3p | 6 | 11 | 0 | 7 | 2 | 0.44444444 | 0 | 0.6520767 |
| rno-miR-146a-5p | 5 | 9 | 11 | 14 | 0.75 | -0.4347826 | -1.1375035 | -0.6374299 |
| rno-miR-369-5p | 5 | 23 | 6 | 12 | 0.18181818 | 0.62857143 | -0.2630344 | 0.93859946 |
| rno-miR-336-5p | 4 | 9 | 13 | 9 | 1.05882353 | 0 | -1.7004397 | 0 |
| rno-miR-211-5p | 4 | 10 | 10 | 12 | 0.85714286 | -0.1818182 | -1.3219281 | -0.2630344 |
| rno-miR-122-5p | 3 | 2 | 25 | 13 | 1.57142857 | -1.4666667 | -3.0588937 | -2.7004397 |
| rno-miR-222-3p | 3 | 39 | 4 | 4 | 0.28571429 | 1.62790698 | -0.4150375 | 3.28540222 |
| rno-miR-411-3p | 1 | 12 | 1 | 6 | 0 | 0.66666667 | 0 | 1 |

